# Cognitive Domains Function Complementation by *NTNG* Gene Paralogs

**DOI:** 10.1101/034645

**Authors:** Pavel Prosselkov, Denis Polygalov, Qi Zhang, Thomas J. McHugh, Shigeyoshi Itohara

## Abstract

Gene duplication was proposed by S.Ohno (1) as a key mechanism of a novel gene function evolution. A pair of gene paralogs, *NTNG1* and *NTNG2*, sharing identical gene and protein structures and encoding similar proteins, forms a functional complement subfunctionalising (SF) within cognitive domains and forming cognitive endophenotypes, as detected by Intellectual Quotient (IQ) tests (2). Both *NTNG* paralogs are associated with autism spectrum disorder (ASD), bipolar disorder (BD) and schizophrenia (SCZ), with unique nonoverlapping segregation among the other 15 cognitive disorders (CD), emphasizing an evolutionary gain-dependent link between advanced cognitive functions and concomitant neurocognitive pathologies. Complementary expression and human brain transcriptome composition of the paralogs explains the observed phenomena of their functional complementarity. The lowest identity among *NTNGs* is found in a middle of encoded by them proteins designated as uknown (Ukd) domain. *NTNG1* contains anthropoid-specific constrained regions, and both genes contain non-coding conserved sequences underwent accelerated evolution in human. *NTNG* paralogs SF perturbates “structure drives function” concept at protein and gene levels. The paralogs function diversification forms a so-called “Cognitive Complement (CC)”, a product of gene duplication and subsequent cognitive subfunction bifurcation among the *NTNG* gene duplicates.

## INTRODUCTION

Complex behaviors arise from a combination of simpler genetic modules that either have evolved separately or co-evolved. Many genes and proteins they encode have been found to be involved in cognitive information processing with a single variant or a single gene generally accounting for only a partial phenotypic variation of a complex trait, such as cognition. Cognition as a quintessence of brain functioning can be viewed as a product of intricately interlinked networks generated by deeply embedded into it gene-nodes with specific or partially overlapping functions. The robustness of the cognitive processing towards its single elements genetic eliminations (to study their function) and its simultaneous fragility expressed in the multiple forms of neurological disorders manifest the existence of cognitive domains interlocked but SF within a unit of cognition formed upon these domains interaction. Previously, we have described a function of a pair of gene paralogs, *NTNG1* and *NTNG2*, involved in human IQ tests performance, and underwent hominin-specific evolutionary changes (2). Hereby, we report on these gene paralogs features focusing on underlying mechanisms of their function segregation and complementation within the cognitive domains.

## RESULTS

The previously observed phenomena of functional complementation among the *NTNG* paralogs within cognitive domains (2) is also manifested in NTNG-associated human pathologies diagnosed in most cases (if only not in all) by a cognitive decline (Figure 1A-1 and A-2). Both genes are associated with BD and SCZ – devastating disorders sharing similar etiology (3) with genetic correlation by multivariate analysis of 0.590 (4), linked to human creativity (5), and characterized by impulsiveness as a common diagnostic feature (6). Recently found associations of both paralogs with ASD (7) supports the reported genetic correlation of 0.194 ASD/SCZ pair (4) and shared module eigengenes detected by PC1 among these two disorders (8). 12 NTNG1-linked CDs, ranging from AD to TS, span a broad spectrum of clinical features frequently involving reduced processing speed (PS) and verbal comprehension (VC, Figure 1A-1). As for *NTNG2*, working memory (WM) deficit and inability “to bind” events (perceptual organization, PO) are the most prominent diagnostic traits for the SLE and TLE patients (Figure 1A-2), with PN also being characterised by indolent behavior in 90% of the cases (9). Interestingly to note that association of the synapse-expressed *NTNG2* with both SCZ and autoimmune pathology (SLE) correlates with a recent finding that human complement component C4 is involved in the synapse elimination and SCZ development (10). Thus, both *NTNG* paralogs are associated with a variety of CDs and mostly in a non-overlapping manner, except for ASD, BD and SCZ characterized by shared and wide spectrum of cognitive abnormalities. The clinical etiology of the aforementioned diseases supports the deduced by IQ functional complementation existing among the *NTNG* paralogs (2) with (VC/PS) and (WM/PO) endophenotypic deficits being uniquely segregated within the associated cognitive pathologies.

**Figure 1.**
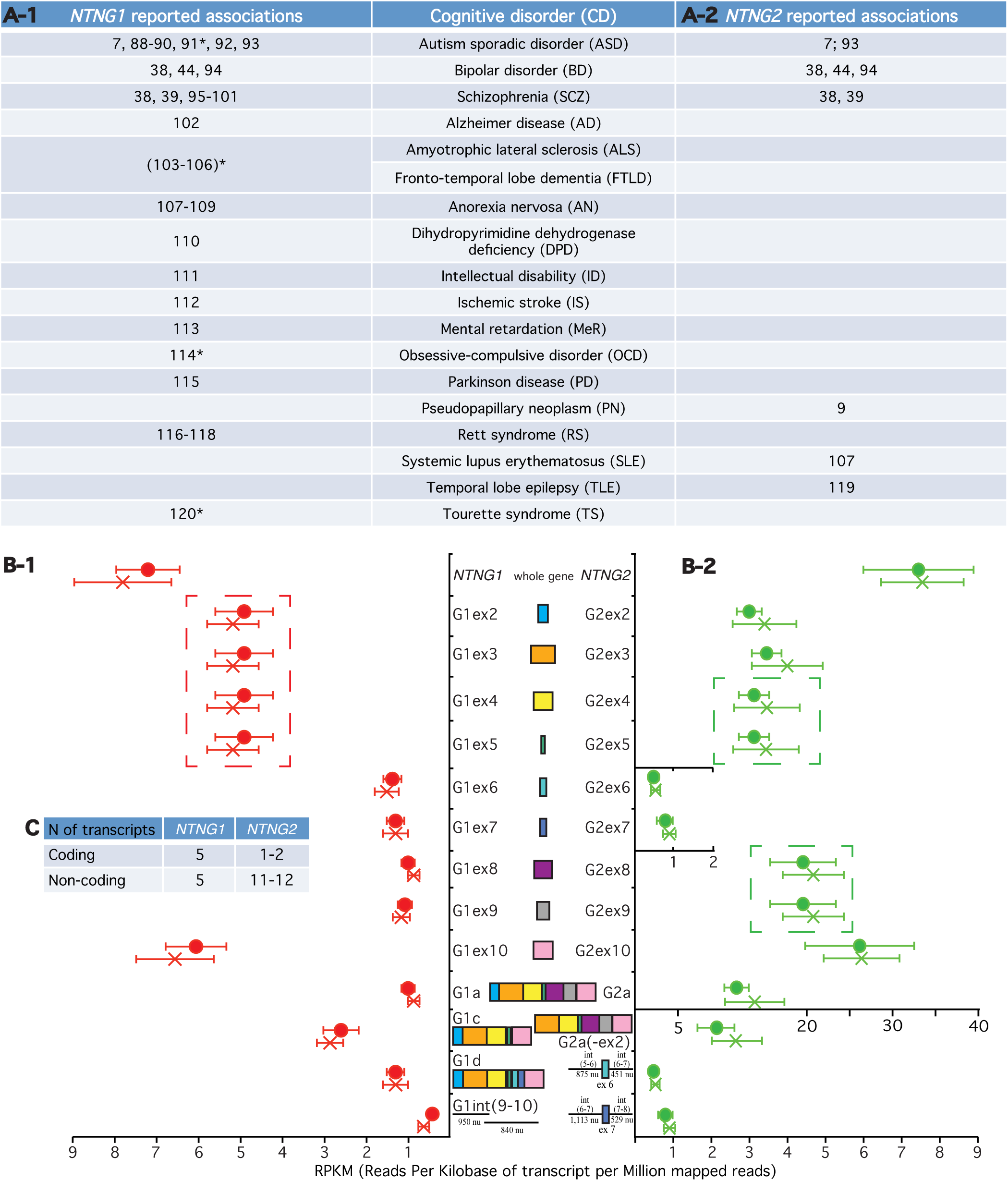
*NTNG* paralogs complementation within neurological disorders and brain transcriptome. (A-1, A-2) Reported cognitive disorders associations for *NTNG1* and *NTNG2*. *denotes rather an indirect association via a direct interaction with the research target. (**B-1, B-2**) RNA-seq of the STG of healthy (circle) and SCZ (cross) human subjects. The original dataset was produced by **Wu et al. (12)**, accession number E-MTAB-1030 on ArrayExpress (**ST1a**) and reprocessed as described in **SM**. Five *NTNG1* and four *NTNG2* transcripts, consistently expressed across all 1 6 human samples are shown. Two samples (one healthy and one SCZ) have been omitted due to unsatisfactory quality of reads and expression profiling (**ST1b**). For the SNPs calling by SAMtools see **ST1c**. Data are presented as a mean RPKMαSEM. (**C**) Total number of the assembled transcripts across all samples for both paralogs (see **ST1d** for the completely reconstructed NTNG transcriptome). Dash-outlined are co-spliced exon clusters.

Since both gene paralogs are expected to have identical gene exon/intron compositions but different in their intron lengths (11) we have reconstructed both paralogs transcriptomes by re-processing the publicly available RNA-seq dataset (12) from healthy and SCZ human subjects superior temporal gyrus (STG) post-mortem brain tissue (Supplementary Table 1a=ST1a). A difference is noted instantly at the total expression levels (genes, exons, individual RNA transcripts) when two gene paralogs are compared (Figure 1B-1 and B-2). *NTNG2* amount (as a whole gene) is 5 times larger comparing to *NTNG1;* exons (2–5) are 3 times, exons (8–9) are 18 times and exon 10 is 4 times higher expressed for *NTNG2* than for *NTNG1*. The only two exons outlaying the prevailing amount rule for the *NTNG2* mRNAs are exons 6 and 7, expressed nearly at the same absolute level as for the *NTNG1* exon paralogs, making them highly underrepresented within the *NTNG2* transcriptome. Next, distinct non-alternating splicing modules are formed by exons (2–5) for *NTNG1* (Figure 1B-1), while exons (4–5) and exons (8–9) for *NTNG2* (Figure 1B-2). Two structurally identical RNA transcript paralogs *(NTNG1a* = G1a and *NTNG2a* = G2a) have been found to exist in both *NTNG* transcriptomes with G2a being expressed at 8-9 times higher level than G1a. *NTNG1* is uniformly presented across the all analysed 16 human samples by 2 more protein coding RNAs (G1c and G1d, detected previously in mice brain, Nakashiba et al., 2000) and by 2 non-coding intron (9–10) derived transcripts (Figure 1B-1). At the same time, *NTNG2* transcriptome is comprised of one extra potentially coding RNA (G2a-like with exon 2 spliced out but in-frame coding preserved) and 2 assumed to be noncoding RNAs with exons 6 and 7 retained along with preceding and following them introns. Quite interesting that these two latter transcripts are the only RNA species with *NTNG2* exon 6 and 7 retained (Figure 1B-2). Two more coding (G1f and G1n) and 4 more non-coding for *NTNG1* and 9 extra non-coding for *NTNG2* RNA species have been also assembled from the available reads but due to inconsistency in their appearance across all 16 STG samples they are not presented on the figure but summarized in the table (Figure 1C, for details refer to ST1d). Summarising above, quantitative and qualitative complementary differences is a prominent feature characterising the brain RNA transcriptome of human *NTNG* paralogs. However, no significant changes at the transcription level of neither whole genes, nor individual exons, nor reconstructed RNA transcripts have been found when SCZ and healthy subjects samples are compared.

Upon calling the presence of IQ-affecting SNPs (2) across all STG samples (ST1c) it has been revealed that 15 out of 16 subjects were positive for the T-allele of rs2149171 (exon 4-nested), shown above to attenuate the WM score in SCZ patients, and making a comparison among the allele carrier *vs* non-carrier impossible. Four healthy and three SCZ samples carry a T-allele of rs3824574 (exon 3-nested, non-affecting IQ), and 1 healthy and 1 SCZ sample each contains a C-allele of rs4915045 (exon10, non-coding part-nested, and non-affecting IQ). Thus, among the eleven cognitive endophenotype-associated SNPs only 3 were possible to call out of the available *NTNG* transcriptome.

Distinctly complementary nature of the *NTNG* paralogs segregation within neurological disorders and RNA transcriptome usage in STG (Figure 1) has prompted us to analyse both genes expression across the entire human brain. We have reconstructed both genes expression profiles in the human brain areas over the life span from conception (pcw = post-conception week) to mature age (30-40 yrs old) using the RNA-seq data from BrainSpan (www.brainspan.org). Similarities and differences are easily noted when the age-dependent phases of *NTNG1* and *NTNG2* expression profiles are matched (Figure 2). Based on the visual inputs three distinct classifiers have been elaborated: 1. predominantly synchronous (Figure 2A(1–4)), characteristic mostly for the cortical areas; 2. predominantly mixed and asynchronous (Figure 2B), characteristic for the cerebellar cortex and subcortical formations; and 3. anti-phasic (complementary, Figure 2C), characteristic for the MD of thalamus and hippocampus. All analysed brain areas demonstrate an elevated level of *NTNG2* expression in comparison to *NTNG1* except for thalamus (Figure 2C) with the largest difference observed at the time of birth (35-37 pcw) or soon after (4 mo) for the synchronous classifiers (Figure 2A), oscillating increment values across the life span for the mixed (Figure 2B) and anti- phasic (Figure 2C) classifiers. It is quite intriguing to note that essentially all brain areas show a trend towards the expression difference being negated between the paralogs by reaching the mature age of 30-40 yrs old (nearly or above the mean age used for the IQ testing), except MD where the expression discrepancy is increased. Thus, the observed functional complementation among the *NTNG* paralogs is supported by the anatomical distribution of the genes in human brain and their expression pattern modality over the human subjects lifetime.

**Figure 2.**
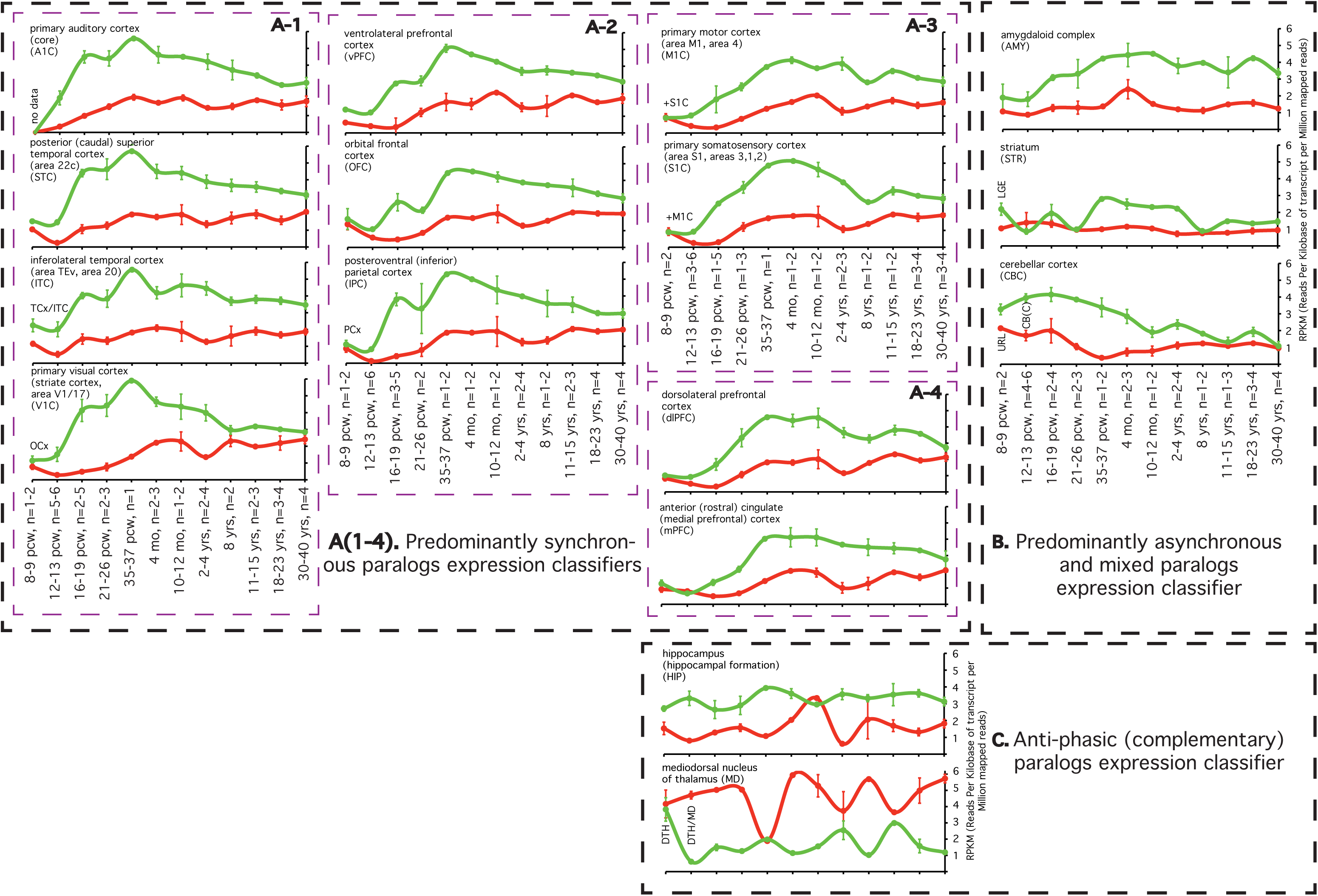
*NTNG* paralogs expression dynamics classification (A-C) in the human brain across the life span. **A (1–4):** further subdivision of the classifier. RNA-seq data are from the BrainSpan (www.brainspan.orq) presented as a meanαSEM. TCx = temporary neocortex; OCx = occipital neocortex; PCx = parietal neocortex; LGE = lateral qanqlionic eminence; CB(C) = cerebellum(cortex); DTH = dorsal thalamus; URL = upper (rostral) rombic limb; pcw = postconception week. Two data points (MD, 12-13 pcw, and mPFC, 16-19 pcw) for *NTNG1* expression were omitted as they were 6-7 times different from the mean of other replicas. All processed brain samples are listed in **ST2**.

A direct comparison of the *NTNG* paralogs shows not only identical intron-exon gene structure (Figure 1B-1, 2B-2) but also closely matched exon sizes (Figure 3A). There are three exons of identical sizes (exons 4, 8 and 9), another three exons differed by one encoded amino acid (exons 3, 5 and 6) and there are exons of different sizes (exons 2, 7 and 10). In terms of size the largest difference among the genes is visually presented by the introns: intron (9–10) of *NTNG1* is 52.7 times larger its *NTNG2* paralogous intron with intron (6–7) of *NTNG1* being only 1.43-times larger pointing towards non-equilibria process of non-coding elements elaborations as the process of gene paralogs SF proceeded. Nevertheless, it can be generalised that in average all *NTNG1* introns are several times larger their *NTNG2* analogs (Figure 3A). We have shown previously that exons 6 and 7 are differentially used within the brain *NTNG* transcriptome (Figure 1B-1 and B-2) and to explore their potential contribution into the paralogs SF we have built identity matrices with these exons being excluded and included (but still producing in-frame existing transcripts, Figure 3B-1 left and right panels, respectively). Exclusion of both exons from the full-lengths transcripts (thus converting *NTNGlm* to *NTNGla* and *NTNG2b* to *NTNG2a*, respectively) increases the identity of DNA on 2% (a relatively large effect since both exons together represent only 7.22 and 9.69% of the total coding part of the full-length RNA transcripts, *NTNG1m* and *NTNG2b*, respectively). This effect becomes even stronger when the encoded by these transcripts proteins are also compared (Figure 3B-2). The spliced out Ukd protein domains (encoded by the exons 6 and 7) increases the proteins identity on 3.8% thus making the middle of both genes (and encoded proteins) substantially more different among the both gene paralogs. To corroborate this observation and to explore the importance of other protein parts we have directly compared the sequences encoded by the full-length transcripts and producing Netrin-G1m and Netrin- G2b (Figure 3C). Similarly to what has been shown on Figure 3B-1 and 3B-2, the lowest identity (17.5%) is represented by the Ukd domain (encoded by the exons 6 and 7) and by the preceding it exon 5 (a 3’-part of the LE1 domain). Two other areas also show a substantially low identity, namely the N-terminus (it includes the protein secretory signal indicated by an arrow) and the outmost C-terminus responsible for the unique feature of Netrin-Gs - the GPI attachment. Thus, based on the percent identity comparisons among the Netrin-G paralogs it can be predicted that there are several potential protein parts contributing to the paralogs SF. As it has been reported by Seiradake et al. (13), identical gene and protein domain compositions result in the identical structural motif with differences only in the spatial arrangement of the loops facing the post-synaptic Netrin-G’s interacting partners, NGL-1 and NGL-2, respectively (Figure 3D). Loop I binding surfaces alignment (Figure 3C, blue color) shows a high level of conservation (with at least 5 amino acids 100% conserved) among the Netrin-G paralogs, indicating that it is unlikely to be responsible for the cognate ligand binding specificity. Neither Loop II (Figure 5C, yellow color) nor Loop III Figure 5C, orange color) display a single conserved amino acid shared among the paralogous binding interfaces (as it has originally been described in (13)). Thus the complementary pattern of the pre-postsynaptic interactions mediated via specific Netrin-G/NGL pairs is reflected in the reciprocally different sizes of the loops binding interfaces representing another element of the NTNG-encoded protein paralogs SF.

**Figure 3.**
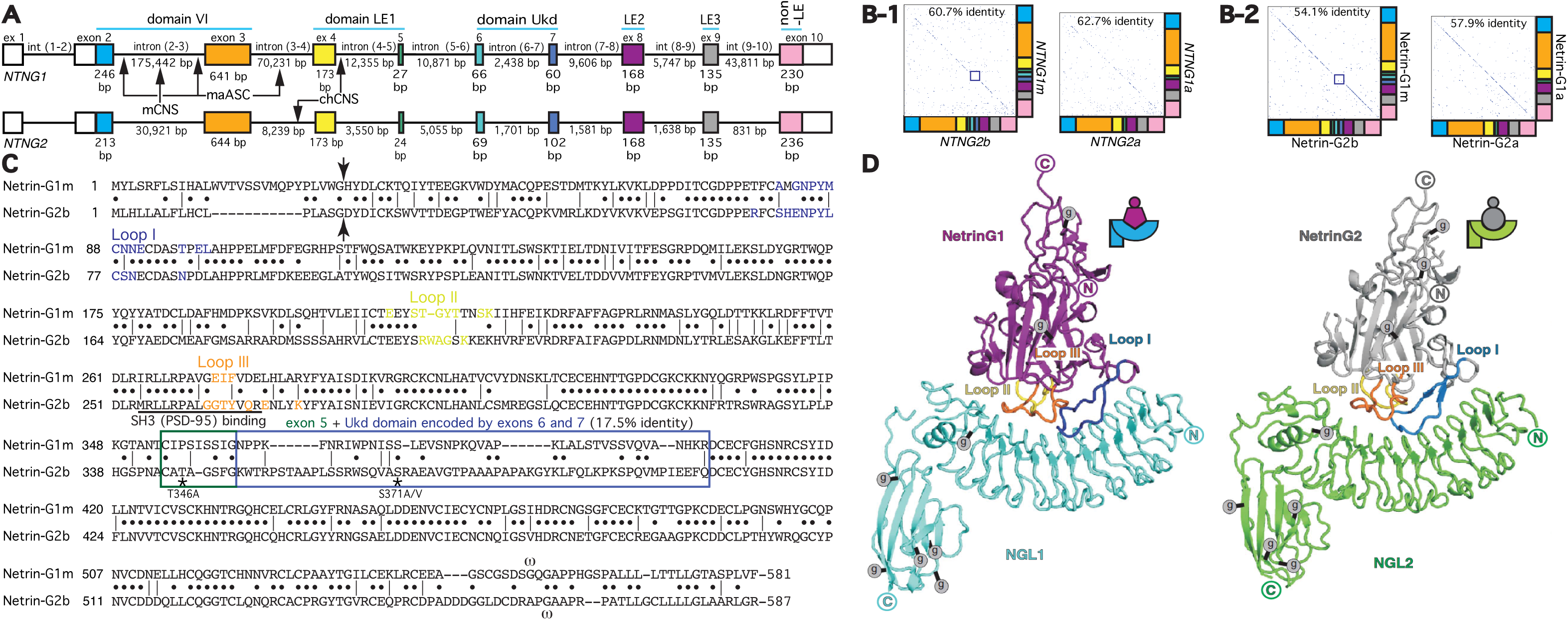
Human *NTNG* paralogs DNA and protein sequence comparisons and “structure-function” rule incongruency. (**A**) Identical gene structures with different sizes of introns. RNA-seq data from Figure 1B were used to precisely deduce the exon/intron junction boundaries. The sizes of exons 1, 10 and introns (1–2) are not indicated due to observed among the splice transcripts lengths variability (see **ST1a** for details). Arrows indicate location of CNS = conserved non-coding sequences underwent accelerated evolution in human compare to mice (mCNS) and chimpanzee (chCNS), as per (**124**); and ASC = anthropoid-specific constrained regions in human compare to marmoset (maASC), as per (**121**). (**B**) Identical exonal composition of the longest *NTNG* encoded RNA paralog transcripts and corresponding proteins with relatively high percent of identity among them dependent on the included/excluded Ukd domain (B-2) encoded by the exons 6 and 7 (**B-1**). Notably, the protein sequence represents higher percent of the paralogs difference than encoded it DNA. The matrices were obtained by GeneJockey II (Biosoft). (C) Protein alignments for the longest human *NTNG* encoded proteins, Netrin- G1m and Netrin-G2b, with Loops I-III highlighting binding sites for their cognate post-synaptic binding partners NGL-1 (*Lrrc4c*) and NGL-2 (*Lrrc4*), respectively, as determined by **Seiradake et al**. (13). Arrow indicates a putative secretory cleavage site location, as calculated by SignalI*p* (122), the blue rectangle delineates the area of the lowest identity (3’-domain LE1+Ukd domain); *m* - denotes a point of putative GPI-attachment, as predicted by Big-PI (123). PSD-95 interaction site via the SH3-binding domain ((**69**), as determined for mice Netrin-G2) overlaps with the Loop III nGl-2 binding surface. Two stars indicate a modern human (T346A) and a hominin-specific (S371A/V) amino acid substitutions (2). (D) Identical structural motif of the Netrin-G1/NGL1 and Netrin-G2/NGL2 complexes as per (13). The figure’s reproduction is covered by the Creative Commons license.

## DISCUSSION

Complementary contribution of *NTNG* paralogs into human cognitive pathologies. Involvement of the pre-synaptically expressed axon-localised *NTNGs* in SCZ diagnosis supports the established view of SCZ as a product of distorted trans-synaptic signaling (14), with a recent study also proving that axonal connectivity-associated genes form a functional network visualisable by fMRI (15), and that brain connectivity predicts the level of fluid intelligence (16, 17). Both *NTNGs* have been found to participate in the brain functional connectivity found by the parcellated connectome reconstruction (18). Most of the reported disease associations link *NTNG1* to SCZ with a variety of other neurologic pathologies (15 in total, Figure 1A-1), while *NTNG2* pathologic associations (6 in total, Figure 1A-2) are quite limited to those affecting WM or PO. Among them is SLE frequently characterized by WM deficit (19) and also known to represent schizoid-type abnormalities characteristic for autoimmune pathologies (20, 21). Immune activation is known to lead to altered pre-pulse inhibition (a key diagnostic trait for SCZ) reversed by antipsychotics (22). The three diseases associated with both paralogs (ASD, BD and SCZ) are also a primary focus of the recently initiated PsychENCODE project (23). It is also worth to mention the resemblance of the reported disease associations with the behavioral phenotypes of *Ntng1* and *Ntng2* gene knockout mice (24).

A gene content associated with attenuated IQ score often relates to numerous diseases, such as SCZ, ASD, depression, and others (see (25) for ref.; 26) with several of them also undergone positive selection following the human brain evolution (27). Despite the fact that global network properties of the brain transcriptome are highly conserved, among the species there are robust human-specific disease-associated modules (28) and human accelerated regions (HARs) - highly conserved parts of genome that underwent accelerated evolution in humans (29). HARs can serve as genomic markers for human-specific traits underlying a recent acquisition of modern human cognitive abilities by brain (30) but that also “might have led to an increase in structural instability... resulted in a higher risk for neurodegeneration in the aging brain” (31), rendering our intellectual abilities genetically fragile (32) and resulting in a variety of CDs. The role genomic context, epistasis (33), plays in the evolution and pathology is manifested by frequently found disease-causing alleles present in animals without obvious pathological symptoms for the host (34). Any CD is characterized by general intellectual disability (GID) plus psychiatric symptoms. A genetic perturbation-exerted behavioral cognitive deficit (BCD) in an animal model organism is a poor match to a human CD *per se* due to very poor contextual resemblance between the human GID and animal BCD together with the absence of interpretable psychiatric symptoms. Usefulness of animals as psychiatric models is also compromised by the fact that transcriptome differences within species tissues is smaller than among the homologous tissues of different species (35, 36). No wonder that the compounds that “cure” mice models consistently fail in human trials (discussed in (37)).

*NTNG* paralogs brain transcriptome intrinsic complementarity and possible mechanism for the IQ-affecting mutation alleles effect. There is no global change at the mRNA level between healthy subjects and SCZ patients (Figure 1B). This conclusion is supported by previously published works stating that globally altered mRNA expression of *NTNG1* or *NTNG2* is unlikely to confer disease susceptibility, at least in the temporal lobe (38), and Brodmann’s area (39). However, the original paper-source of the STG samples RNA-seq along with many other genes (>1,000) found that *NTNG1* (but not *NTNG2)* falls within the group of genes with significant alternative promoter usage ((12): ST6, *p*<9.05E-10 at FDR < 0.5) and *NTNG2* (but not *NTNG1)* clusters with genes (>700) with significant alternative splicing change ((12): ST7, *p*<6.15E-12 at FDR 0.5) when SCZ and controls are compared. Such GWAS observation adds an extra layer of complementary regulation to both *NTNG* paralogs on a top of the described in the results section complementary usage rule for the exons, formed by unspliced splicing modules, resulting transcripts and comprising them exons (Figure IB). Based on the available RNA-seq dataset it was almost impossible to detect RNA with the matched position of *NTNG* SNPs used for the IQ testing (ST2c) except for two coding exons located (rs2149171 and rs3824574) and exon 10 non-coding area located but transcribed rs4915045 (in 2 out of 16 samples). This fact points towards indirect effect of the IQ-affecting mutation alleles potentially associated with shorter (secretable) isoforms generation (Prosselkov et al., unpublished) lacking two of the most prominent *NTNG* features: GPI-link and the Ukd domain through an aberrant splicing factor binding. The GPI-link is a hallmark of Netrin-G family members (40, 41) and lacking it the aberrant Netrin-G isoforms are likely to mimic the action of their releasable ancestry molecules - netrins, still being able to bind to their cognate postsynaptic ligand - NGL but without forming an axonal-postsynaptic contact and potentially dominant negative consequences. The Ukd domain of Netrin-G1, despite its so-far unknown function, is involved in lateral binding to the pre-synaptically localised LAR modulating the binding strength between NGL-1 and Netrin-G1 (42). Work is currently underway in search for a similar lateral interaction partner for the Netrin-G2 Ukd domain. Inclusion of the Ukd-encoding domain exons 6 and 7 is regulated by the Nova splicing factor (43) affecting the cortex Netrin-G1 exon 7 but not exon 6, and, simultaneously, Netrin-G2 paralog exons exhibiting an opposite pattern. In general, it is tempting to speculate that deregulation of *NTNG* transcripts processing may have a role in the brain-controlled cognitive abilities and associated CDs. Supporting such notion, a decreased level of Netrin-G1c mRNA (exons 6-9 excluded, Figure 1B-1) has been reported for BD and SCZ (38) with Netrin-G1d (exons 6 and 7 included but 8-9 excluded, Figure 1B-1) and Netrin-G1f (a secretable short isoform consisted of domain VI only and lacking the Ukd and GPI-link) being increased in BD, but not in SCZ, in anterior cingulated cortex (44).

Higher Netrin-G1d mRNA expression in fetal brain but low for the Netrin-G1c isoform in the human adult (38) indicates different functionality of these two splice variants joggling with the Ukd domain inclusion/exclusion pattern. And, according to our unpublished data, if Netrin-G1 Ukd-containing isoforms are the dominant isoforms in adult mouse brain, Netrin- G2 Ukd-containing isoforms are present only at the trace level (Prosselkov et al., forthcoming), resembling the human STG samples transcriptome pattern (Figure 1B-1 and B-2). A similar “dynamic microexon regulation” associated with the protein interactome misregulation has been reported to be linked to ASD (45).

Synchronous and complementary expression of *NTNG* paralogs in the human brain supports the IQ-associated cognitive endophenotypes. Influential parieto-frontal integration theory (P- FIT, (46)) states that general intelligence (“g”) is dependent on multiple brain cortical areas such as dlPFC, Broca's and Wernicke's areas, somatosensory and visual cortices (47). Despite “g” is widely accepted as the only correlate of the intelligence, its unitary nature was challenged by (48) claiming had indentified two independent brain networks (for memory and for reasoning) responsible for the task performance, the idea later criticised for the employed data processing approach (49). Higher IQ scores (a composite surrogate of “g”) have been reportedly associated with the fronto-parietal network (FPN) connectivity (50, 51). Higher levels of *NTNG* paralogs expression within the cognition intensively loaded areas of the brain and the distinct patterns of expression profiles (synchronous, asynchronous/mixed, and complementary, Figure 2A) support associations of *NTNG1* and *NTNG2* with the recorded cognitive endophenotypes (2). Based on the expression patterning over the human life-span, among the total 16 analysed brain areas we found two falling under the same “anti- phasic (complementary)” classifier (Figure 2C): HIP and MD. Adding more to that, MD is the only brain area (out of the 16 presented) where *NTNG1* expression level exceeds that of *NTNG2* making it a promising candidate for the phenomena of *NTNGs* SF explanation. Two other brain areas classified by a synchronous paralogs expression deserve a special attention, dlPFC and mPFC (Figure 2A-4). PFC circuitry has been known as a “hub of the brain’s WM system” (52, 53), which acts through direct HIP afferents (54) and has many connections with other cortical and subcortical areas (55). mPFC may function as an intelligence-control switchboard and lPFC, part of the FPN global connectivity, predicts the WM performance and fluid intelligence (56). Interactions of the auditory recognition information fed by the vPFC stream with the sequence processing by the dorsal stream are crucial for the human language articulation (57; 58). The fact that both *NTNG* paralogs are extensively expressed across PFC (Figure 2A-2 and A-4) pinpoints this area as a key for future molecular studies of *NTNGs* and the human-unique symbolic communications. And PFC is not only implicated in many psychiatric disorders, including SCZ ((59); see also (55) for ref.), but is also the only brain structure unique to primates without known homologs in the animal kingdom (60). Evolution of the protein paralogs encoded by the *NTNGs*. Forkhead box P2 (FOXP2) - a ubiquitously expressed transcription factor that has been reportedly linked to the evolution of human language through T303N, N325S substitutions when compared to a primate ortholog (61), is 100% identical to Nea protein (62). FOXP2 regulates expression of multiple genes in human and chimpanzee (63), and among them is an M3 gene brain module representative responsible for general fluid cognitive abilities (26), *LRRC4C*, a gene encoding NGL-1, a post-synaptic target of Netrin-G1. Similarly to FOXP2, Netrin-G1 is a 100% conserved protein among the hominins with only 1 mutation found in chimpanzee which is absent in marmoset (and other primates) and mice proteins (2). On the other hand, extinct hominins’ Netrin-G2 relative to modern human contains T346A point mutation (as per current version of hg19), also found in primates and mouse and known as rs4962173 (dbSNP missense mutation) representing an ancient substitution from Neandethal genomes found in modern humans and reflecting a recent acquisition of the novel allele around 5,300 yrs BC. Nothing is known regarding the functional significance of this mutation allele but biochemically a substitution of alanine (A) on a polar threonine (T) could bring an extra point of post- translational modification, e.g. a phosphorylation or glycosylation (NetPhos2.0 (64) assigns a low probability score for the T346 to be phosphorylated but Net0Glyc4.0 (65) robustly predicts it to be glycosylated, see SM). Another mutation, S371A/V, reflects a selective sweep in Netrin-G2 protein from primates to hominins within a similar to T346A functional context when a hydrophobic alanine (in chimpanzee, A)/valine (in marmoset, V) is replaced by a polar serine (S) with a strong positive predictions for glycosylation but not phosphorylation (see SM). This poses a question whether these two human-specific protein substitutions associate with advanced cognitive traits as they may represent a hidden layer of poorly studied so far protein glycosylation-associated regulatome known to affect the brain function and diseases (66, 67). Adding more to this, T346 is nested on exon 5 just 20 nu away from the affecting WM score rs2274855 mutation allele (2), and, together with S371A/V, they are both located within the lowest percent identity area (exons (5–7)) of Netrin-Gs (Figure 3C) and, proposedly, contributing to the *NTNG* duplicates SF. There are at least three more protein parts potentially contributing to the gene paralogs specialised function subdivision (based on the low identity scores, Figure 3C): the secretory peptide, the GPI-link, and the outmost structurally elaborated unstructured loops (I-III) responsible for the reciprocal binding of Netrin-Gs to their post-synaptic cognate partners, NGL-1 or NGL-2, both containing a C-terminal PDZ-binding domain (68). An interesting finding was reported in (69) reported a presence of SH3(PSD95) domain binding site (required for the phosphatidylinositol-3-kinase recruitment) in mice Netrin-G2 (100% identical to human) but not in Netrin-G1. The detected SH3 binding site overlaps with the Netrin-G2-loop III responsible for the binding specificity to NGL-2 (12, 70, 71). A plausible working hypothesis would be that while internalised (and being GPI-link naive/immature) the pre-synaptic

Netrin-G2 is bound to SH3-PSD95 via loop III but as soon as being secreted extracellularly (and being attached to the membrane) it is bound to post-synaptic NGL-2. Corroborating this, in the absence of Netrin-G2 in the KO mice NGL-2 is unstable on the post-synaptic surface and gets quickly internalised (24). We can only speculate regarding the potential importance of PSD-95(SH3)-Netrin-G2-NGL-2 scaffolding loop interaction/competition but the ability for Netrin-G1 to bind to SH3 has not been reported. Following this logic, Netrin-G1 should have a similar binding partner via loop II.

The overall identical structural scaffold among the Netrin-G paralogs (Figure 3D) is likely to represent an anciently preserved one of the primordial protein (encoded by a single gene in the primitive urochordate *C.intestinalis*) and its contribution to the process of SF among the *NTNG* paralogs goes against the “structure drives function” concept. It looks like that it is not the “structure” but rather the “evolution” itself that drives a selection for the best structural (or unstructural in our case) fit out of the available frameworks provided by the gene duplicates to fulfill the emerged functional demand in a new ecological niche. The intricate variability of phenotype is grounded by the conserved nature of genotype and constrained by the “structure-function” limitations of the coding DNA and is only possible due to permissive evolutionary continuing elaborations of non-coding areas able to absorb the most recently acquired elements (having a potential to become regulatory at some point, e.g. like HAR5 (30)) and carried over by the neutral drift. At the same time, the multiple protein substitutions coinciding with the SF labor segregation phenomena among the Netrin-G paralogs question their neutral nature. Both of them undergo a purifying selection from mice to human through the reduction in size of non-coding DNA (introns) and encoded proteins (the mice Netrin-G2 is 2 aa longer its human ortholog) further contributing to the host- specific SF. Thus while the non-coding sequences are used to explore the evolutionary space in time, the restrictive boundaries of the paralogs SF are determined by the protein (unstructured) elements.

Molecular evolution of the Cognitive Complement (CC). Appearance of the neural crest (72). an event that “affected the chordate evolution in the unprecedented manner” (73), multipotenl progenitor cells (74), and neurogenic placodes (suggesting a chemosensory and neurosecretory activities (75)) in the first primitive urochordates/tunicates coincides with the presence of *Ntng* precursor gene (ENSCING00000024925) later undergoing two rounds oi duplication events in fish and found to affect human cognitive abilities (2). *NTNG* paralogs are expressed in the human neural crest-forming cells with *NTNG2* 10 times stronger than *NTNG1* (76), both are differentially expressed in human comparing to chimpanzee and rhesus monkey with *NTNG2* expression model showing stronger probability than *NTNG1* (77), and both are stronger expressed in human telencephalon comparing to chimpanzee and macaque (78). *NTNG1* has been classified as a brain module hub gene “whose pattern fundamentally shifted between species” (18). Belonging to distinct modules of brain expression regulation (78, 79), *NTNGs* are classified as “genes with human-specific expression profiles” (79). The nearby gene ~260 kbp upstream of *NTNG2* is *MED27* (mediator of RNA polymerase II) has been proposed to be associated with the evolution of human-specific traits (80). *NTNG1* has also been reported among the “adaptive plasticity genes” (81) potentiating rapid adaptive evolution in guppies *(NTNG2* was not found in the input RNA).

Complementarity among the *NTNG* paralogs and encoded by them proteins has been reported previously: brain expression complementary pattern (in almost self-exclusive manner) defined by the 5’-UTR-localised c/s-regulatory elements (82); complementary distribution within the hippocampal laminar structures (83); axon-dendrite synaptic ending resulting in differential control over the neuronal circuit plasticity (84); mutually-exclusive binding pattern to post-synaptic partners, NGL-1 and NGL-2, dictated by the protein unstructural elements (13); alternative promoter usage *vs* alternative mRNA splicing (12) and increased coefficient of variation (CV, ST1d) for *NTNG1* expression but not *NTNG2* in SCZ patients (also reported in (85)); KO mice behavioral phenotypes and subcellular signaling partners complementarity (24); “differential stability” brain modules expression *(NTNG1* is expressed in the dorsal thalamus (M11) as a hub gene (Pearson’s 0.92) while *NTNG2* is in neocortex and claustrum module (M6, Pearson’s 0.65)) (18); hypocretin neurons-specific expression of *NTNG1* (but not *NTNG2*) as a sleep modulator (86); top-down *vs* bottom-up information flows gating in mice and differential modality (Prosselkov et al., forthcoming), and human IQ-compiling cognitive domains complementation (2). The current study reports on *NTNGs* complementarity association with the CDs (Figure 1A); mRNA splicing pattern complementary at the quantitative and qualitative levels via differential use of the middle- located exons (Figure 1B); brain complementary oscillatory expression over the human life span observed in the intensive cognitively loaded brain areas (Figure 2); AE of the paralogs- segregated unique non-coding elements (Figure 3A); complementary pattern of the protein orthologs (mice-to-human) protein sequence evolution. Such multi-level complementation is likely to reflect a shared evolutionary origin from a single gene in a primitive vertebrate organism 700 mln yrs ago and its subsequent functional segregation among the evolution-generated gene duplicates in jawless fish, such as lamprey.

Occupying independent but intercalating functional niches, *NTNG1* and *NTNG2* do not compensate but complement each other’s function forming a “functional complement” of genes. Half a billion yrs ago the doubled gene dosage led to the gradual SF and manifested in a function complementation within the cognitive domains, at least in human. We would like to coin such gene pair as a Cognitive Complement (CC).

## CONCLUSION

The emerged functional redundancy, as an outcome of gene duplication, leads to function subdivision and its bifurcation among the gene paralogs resulting in the paralogs SF. A functional compensation is known to exist among the evolutionary unrelated genes but has not been reported among the gene paralogs, more frequently characterized by the function complementation. Gene paralogs structural identity (at both, gene and protein levels) does not provide a substrate for function compensation but rather for complementation, perturbating “structure drives function” rule. A gene duplication event of a tunicate *NTNG* primordial gene and the subsequent process of its function specialisation (driven by the new ecological niches appearance and evolution) among the gene duplicates made them to SF into distinct cognitive domains in a complementary manner forming a CC. In our forthcoming work we are to describe how *Ntng* mice genes function resembles that of human orthologs (Prosselkov et al., forthcoming).

## MATERIALS AND METHODS

Human brain *NTNG* transcriptome reconstruction. Relates to Figure 1B and 1-C. The original source of the dataset was produced by ((12): E-MATB-1030) and the downloaded .bam files used for the re-processing are listed in ST1a. All reconstructed transcripts are presented in ST1d standalone Excel file. Two samples were excluded from the analysis due to failed “per base sequence quality” measure, and zero expression level for *NTNG1a* and *NTNG1int(9–10)* otherwise consistently expressed throughout other samples (ST1b). SAMtools software was used for the SNPs calling from the available RNA-seq datasets (ST1c). For details refer to SM.

Human brain expression profiling for *NTNGs* across the life span. The original source of data was www.brainspan.org. All available samples were initially included into the analysis but two of them excluded at a later stage (MD for 12-13 pcw and mPFC for 16-19 pcw) due to high deviation (6-7 times) from the mean for other replicas. The mean expression values per each brain area as RPKM were plotted against the sampling age. Profiles classification was done visually considering the trend over the all plotted points as an average.

*NTNG1 (NTNG1m*) and *NTNG2 (NTNG2b*) full-length mRNA transcripts assembly. Relates to Figure 3B. Human *NTNG1m* brain transcript has been reported previously (125) and we have also confirmed its ortholog presence in the mice brain via full-length cloning (42). Since NCBI contains only its partial CDS (AY764265), we used the RNA-seq-generated exons (Figure 1B) to reconstruct its full-length and to generate an ORF of the encoded Netrin-G1m. Similarly, human *NTNG2b* was reconstructed from the RNA-seq dataset and from Ensemble as follows. Exon 5 sequence was deduced from ENST00000372179, other exons were from ENST00000467453 (no longer available on the current version of Ensemble) except for exon 6 deduced by running three independent alignments against the human genomic DNA with the mice 3’-intron (5–6), exon 6, and 5’-intron (6–7) concomitantly confirmed by the generated full-length ORF for Netrin-G2b. The reconstructed protein was predicted to encode 587 amino acids, which is in a close proximity to the mice netrin-G2b ortholog of 589 residues (Prosselkov et al., forthcoming).

Full-lengths gene structures of *NTNG* paralogs reconstruction. Relates to Figure 3A. Both, the obtained above from the STG brain samples RNA-seq and the reconstructed full-lengths transcripts carrying all stably expressed exons were used to confirm the intron-exon junctions positioning for *NTNG1* and *NTNG2*. Due to observed variability in the intron (1–2) and exon 10 sizes their boundaries were left unmarked.

## SUPPLEMENTARY MATERIALS (SM)

Contain Supplementary Methods (RNA-seq of STG re-processing and SNPs detection) and Supplementary Tables (ST1a-d, ST2) as a single compiled pdf file. Reconstructed RNA-seq (.gtf) of the STG is presented as a standalone Excel file (ST1d). Also included: Netrin-G2b predicted phosphorylation and *O*-glycosylation, Netrin-G1 *vs* Netrin-G2 Ukd alignment (87), predicted secretory peptide cleavage and GPI attachments.

## ACKNOWLEDGEMENTS

This work was in part supported by the “Funding Program for World-Leading Innovative R&D on Science and Technology (FIRST Program)” initiated by the Council for Science and Technology Policy (CSTP), and KAKENHI 15H04290 from the Japan Society for the Promotion of Science (JSPS).

## COMPETING INTERESTS

Authors would like to express a lack of any competing interests associated with the work.

## REFERENCES

1. Ohno S. 1970. Evolution by Gene Duplication. Springer, New York.

2. Prosselkov P, Hashimoto R, Polygalov D, Kazutaka O, Zhang Q, McHugh TJ, Takeda M, Itohara S. 2015. Cognitive Endophenotypes of modern and extinct hominins associated with NTNG gene paralogs. BioRxiv, doi: 10.1101/034413.

3. Lee SH, Ripke S, Neale BM, Faraone SV, Purcell SM, Perlis RH, Mowry BJ, Thapar A, Goddard ME, Witte JS, Absher D, Agartz I, Akil H, Amin F, Andreassen OA, Anjorin A, Anney R, Anttila V, Arking DE, Asherson P, Azevedo MH, Backlund L, Badner JA, Bailey AJ, Banaschewski T, Barchas JD, Barnes MR, Barrett TB, Bass N, Battaglia A, Bauer M, Bayés M, Bellivier F, Bergen SE, Berrettini W, Betancur C, Bettecken T, Biederman J, Binder EB, Black DW, Blackwood DHR, Bloss CS, Boehnke M, Boomsma DI, Breen G, Breuer R, Bruggeman R, Cormican P, Buccola NG, Buitelaar JK, Bunney WE, Buxbaum JD, Byerley WF, Byrne EM, Caesar S, Cahn W, Cantor RM, Casas M, Chakravarti A, Chambert K, Choudhury K, Cichon S, Cloninger CR, Collier DA, Cook EH, Coon H, Cormand B, Corvin A, Coryell WH, Craig DW, Craig IW, Crosbie J, Cuccaro ML, Curtis D, Czamara D, Datta S, Dawson G, Day R, De Geus EJ, Degenhardt F, Djurovic S, Donohoe GJ, Doyle AE, Duan JB, Dudbridge F, Duketis E, Ebstein RP, Edenberg HJ, Elia J, Ennis S, Etain B, Fanous A, Farmer AE, Ferrier IN, Flickinger M, Fombonne E, Foroud T, Frank J, Franke B, Fraser C, Freedman R, Freimer NB, Freitag CM, Friedl M, Frisén L, Gallagher L, Gejman PV, Georgieva L, Gershon ES, Geschwind DH, Giegling I, Gill M, Gordon SD, Gordon-Smith K, Green EK, Greenwood TA, Grice DE, Gross M, Grozeva D, Guan WH, Gurling H, De Haan L, Haines JL, Hakonarson H, Hallmayer J, Hamilton SP, Hamshere ML, Hansen TF, Hartmann AM, Hautzinger M, Heath AC, Henders AK, Herms S, Hickie IB, Hipolito M, Hoefels S, Holmans PA, Holsboer F, Hoogendijk WJ, Hottenga JJ, Hultman CM, Hus V, Ingason A, Ising M, Jamain S, Jones EG, Jones I, Jones L, Tzeng JY, Kähler AK, Kahn RS, Kandaswamy R, Keller MC, Kennedy JL, Kenny E, Kent L, Kim Y, Kirov GK, Klauck SM, Klei L, Knowles JA, Kohli MA, Koller DL, Konte B, Korszun A, Krabbendam L, Krasucki R, Kuntsi J, Kwan P, Landen M, Längström N, Lathrop M, Lawrence J, Lawson WB, Leboyer M, Ledbetter DH, Lee PH, Lencz T, Lesch KP, Levinson DF, Lewis CM, Li J, Lichtenstein P, Lieberman JA, Lin DY, Linszen DH, Liu CY, Lohoff FW, Loo SK, Lord C, Lowe JK, Lucae S, MacIntyre DJ, Madden PAF, Maestrini E, Magnusson PKE, Mahon PB, Maier W, Malhotra AK, Mane SM, Martin CL, Martin NG, Mattheisen M, Matthews K, Mattingsdal M, McCarroll SA, McGhee KA, McGough JJ, McGrath PJ, McGuffin P, McInnis MG, McIntosh A, McKinney R, McLean AW, McMahon FJ, McMahon WM, McQuillin A, Medeiros H, Medland SE, Meier S, Melle I, Meng F, Meyer J, Middeldorp CM, Middleton L, Milanova V, Miranda A, Monaco AP, Montgomery GW, Moran JL, Moreno-De-Luca D, Morken G, Morris DW, Morrow EM, Moskvina V, Muglia P, Mühleisen TW, Muir WJ, Muller-Myhsok B, Murtha M, Myers RM, Myin-Germeys I, Neale MC, Nelson SF, Nievergelt CM, Nikolov I, Nimgaonkar V, Nolen WA, Nöthen MM, Nurnberger JI, Nwulia EA, Nyholt DR, O’Dushlaine C, Oades RD, Olincy A, Oliveira G, Olsen L, Ophoff RA, Osby U, Owen MJ, Palotie A, Parr JR, Paterson AD, Pato CN, Pato MT, Penninx BW, Pergadia ML, Pericak-Vance MA, Pickard BS, Pimm J, Piven J, Posthuma D, Potash JB, Poustka F, Propping P, Puri V, Quested DJ, Quinn EM, Ramos-Quiroga JA, Rasmussen HB, Raychaudhuri S, Rehnström K, Reif A, Ribases M, Rice JP, Rietschel M, Roeder K, Roeyers H, Rossin L, Rothenberger A, Rouleau G, Ruderfer D, Rujescu D, Sanders AR, Sanders SJ, Santangelo SL, Sergeant JA, Schachar R, Schalling M, Schatzberg AF, Scheftner WA, Schellenberg GD, Scherer SW, Schork NJ, Schulze TG, Schumacher J, Schwarz M, Scolnick E, Scott LJ, Shi JX, Shilling PD, Shyn SI, Silverman JM, Slager SL, Smalley SL, Smit JH, Smith EN, Sonuga-Barke EJS, St Clair D, State M, Steffens M, Steinhausen HC, Strauss JS, Strohmaier J, Stroup TS, Sutcliffe JS, Szatmari P, Szelinger S, Thirumalai S, Thompson RC, Todorov AA, Tozzi F, Treutlein J, Uhr M, van den Oord EJCG, Van Grootheest G, Van Os J, Vicente AM, Vieland VJ, Vincent JB, Visscher PM, Walsh CA, Wassink TH, Watson SJ, Weissman MM, Werge T, Wienker TF, Wijsman EM, Willemsen G, Williams N, Willsey AJ, Witt SH, Xu W, Young AH, Yu TW, Zammit S, Zandi PP, Zhang P, Zitman FG, Zöllner S, International Inflammatory Bowel Disease Genetics Consortium (IIBDGC), Devlin B, Kelsoe JR, Sklar P, Daly MJ, O'Donovan MC, Craddock N, Sullivan PF, Smoller JW, Kendler KS, Wray NR. 2013. Genetic relationship between five psychiatric disorders estimated from genome-wide SNPs. Nature Genetics 45:984–994. doi: 10.1038/ng.2711.

4. Maier R, Moser G, Chen G-B, Ripke S, Cross-disorder Working Group of the Psychiatric Genomics Consortium, Coryell W, Potash JB, Scheftner WA, Shi J, Weissman MM, Hultman CM, Landen M, Levinson DF, Kendler KS, Smoller JW, Wray NR, Lee SH. 2015. Joint Analysis of Psychiatric Disorders Increases Accuracy of Risk Prediction for Schizophrenia, Bipolar Disorder, and Major Depressive Disorder. The American Journal of Human Genetics 96:283–294. doi: 10.1016/j.ajhg.2014.12.006.

5. Power RA, Steinberg S, Bjornsdottir G, Rietveld CA, Abdellaoui A, Nivard MM, Johannesson M, Galesloot TE, Hottenga JJ, Willemsen G, Cesarini D, Benjamin DJ, Magnusson PKE, Ullén F, Tiemeier H, Hofman A, van Rooij FJA, Walters GB, Sigurdsson E, Thorgeirsson TE, Ingason A, Helgason A, Kong A, Kiemeney LA, Koellinger P, Boomsma DI, Gudbjartsson D, Stefansson H, Stefansson K. 2015. Polygenic risk scores for schizophrenia and bipolar disorder predict creativity. Nature Neuroscience 18:953–955. doi: 10.1038/nn.4040.

6. Reddy LF, Lee J, Davis MC, Altshuler L, Glahn DC, Miklowitz DJ, Green MF. 2014. Impulsivity and Risk Taking in Bipolar Disorder and Schizophrenia. Neuropsychopharmacology 39:456–463. doi: 10.1038/npp.2013.218.

7. Sanders SJ, He X, Willsey AJ, Ercan-Sencicek AG, Samocha KE, Cicek AE, Murtha MT, Bal VH, Bishop SL, Dong S, Goldberg AP, Jinlu C, Keaney III JF, Klei L, Mandell JD, Moreno-De-Luca D, Poultney CS, Robinson EB, Smith L, Solli-Nowlan T, Su MY, Teran NA, Walker MF, Werling DM, Beaudet AL, Cantor RM, Fombonne E, Geschwind DH, Grice DE, Lord C, Lowe JK, Mane SM, Martin DM, Morrow EM, Talkowski ME, Sutcliffe JS, Walsh CA, Yu TW, Autism Sequencing Consortium, Ledbetter DH, Martin CL, Cook EH, Buxbaum JD, Daly MJ, Devlin B, Roeder K, State MW. 2015. Insights into Autism Spectrum Disorder Genomic Architecture and Biology from 71 Risk Loci. Cell 87:1215–1233. doi: 10.1016/j.neuron.2015.09.016.

8. Parikshak NN, Gandal MJ, Geschwind DH. 2015. Systems biology and gene networks in neurodevelopmental and neurodegenerative disorders. Nature Reviews Genetics 16:441–458. doi: 10.1038/nrg3934.

9. Cavard C, Audebourg A, Letourneur F, Audard V, Beuvon F, Cagnard N, Radenen, B, Varlet P, Vacher-Lavenu M-C, Perret C, Terris B. 2009. Gene expression profiling provides insights into the pathways involved in solid pseudopapillary neoplasm of the pancreas. Journal of Pathology 218:201–209. doi: 10.1002/path.2524.

10. Sekar A, Bialas AR, de Rivera H, Davis A, Hammond TR, Kamitaki N, Tooley K, Presumey J, Baum M, Van Doren V, Genovese G, Rose SA, Handsaker RE, Schizophrenia Working Group of the Psychiatric Genimics Consortium, Daly MJ, Carroll MC, Stevens B, McCarroll SA. 2016. Schizophrenia risk from complex variation of complement component 4. Nature, advance online. doi: 10.1038/nature16549.

11. Yin Y, Miner JH, Sanes JR. 2002. Laminets: Laminin- and netrin-related genes expressed in distinct neuronal subsets. Molecular and Cellular Neuroscience 19:344–358. doi: 10.1006/mcne.2001.1089.

12. Wu JQ, Wang X, Beveridge NJ, Tooney PA, Scott RJ, Carr VJ, Cairns MJ. 2012. Transcriptome Sequencing Revealed Significant Alteration of Cortical Promoter Usage and Splicing in Schizophrenia. PLoS One 7:e36351. doi: 10.1371/journal.pone.0036351.

13. Seiradake E, Coles CH, Perestenko PV, Harlos K, McIlhinney RAJ, Aricescu AR, Jones EY. 2011. Structural basis for cell surface patterning through NetrinG-NGL interactions. The EMBO Journal 30:4479–4488. doi: 10.1038/emboj.2011.346.

14. Lips ES, Cornelisse LN, Toonen RF, Min JL, Hultman CM, the International Schizophrenia Consortium, Holmans PA, O’Donovan MC, Purcell SM, Smit AB, Verhage M, Sullivan PF, Visscher PM, Posthuma D. 2012. Functional gene group analysis identifies synaptic gene groups as risk factor for schizophrenia. Molecular Psychiatry 17:996–1006. doi: 10.1038/mp.2012.37.

15. Richiardi J, Altmann A, Milazzo A-C, Chang C, Chakravarty MM, Banaschewski T, Barker GJ, Bokde ALW, Bromberg U, Büchel C, Conrod P, Fauth-Bühler M, Flor H, Frouin V, Gallinat J, Garavan H, Gowland P, Heinz A, Lemaitre H, Mann KF, Martinot J-L, Nees F, Paus T, Pausova Z, Rietschel M, Robbins TW, Smolka MN, Spanagel R, Ströhle A, Schumann G, Hawrylycz M, Poline J-B, Greicius MD, IMAGEN consortium. 2015. Correlated gene expression supports synchronous activity in brain networks. Science 348:1241–1244. doi: 10.1126/science.1255905.

16. Finn ES, Shen X, Scheinost D, Rosenberg MD, Huang J, Chun MM, Papademetris X, Constable RT. 2015. Functional connectome fingerprinting: identifying individuals using patterns of brain connectivity. Nature Neuroscience online. doi:10.1038/nn.4135.

17. Pamplona GSP, Neto GSS, Rosset SRE, Rogers BP, Salmon CEG. 2015. Analyzing the association between functional connectivity of the brain and intellectual performance. Frontiers in Human Neuroscience 9:A61. doi: 10.3389/fnhum.2015.00061.

18. Hawrylycz M, Miller JA, Menon V, Feng D, Dolbeare T, Guillozet-Bongaarts AL, Jegga AG, Aronow BJ, Lee C-K, Bernard A, Glasser MF, Dierker DL, Menche J, Szafer A, Collman F, Grange P, Berman KA, Mihalas S, Yao Z, Stewart L, Barabasi A-L, Schulkin J, Phillips J, Ng L, Dang C, Haynor DR, Jones A, Van Essen DC, Koch C, Lein E. 2015. Canonical genetic signatures of the adult human brain. Nature Neuroscience 18:1832–1844. doi: 10.1038/nn.4171.

19. Shucard JL, Lee WH, Safford AS, Shucard DW. 2011. The Relationship Between Processing Speed and Working Memory Demand in Systemic Lupus Erythematosus: Evidence From a Visual N-Back Task. Neuropsychology 25:45–52. doi: 10.1037/a0021218.

20. Guilloux JP, Gaiteri C, Sibille E. 2010. Network analysis of positional candidate genes of schizophrenia highlights ... more than ... myelin-related pathways. Molecular Psychiatry 15:786–788. doi: 10.1038/mp.2009.68.

21. Eaton WW, Byrne M, Ewald H, Mors O, Chen CY, Agerbo E, Mortensen PB. 2006. Association of schizophrenia and autoimmune diseases: Linkage of Danish National Registers. American Journal of Psychiatry 163:521–528. doi: 10.1176/appi.ajp.163.3.521.

22. Romero E, Ali C, Molina-Holgado E, Castellano B, Guaza C, Borrell J. 2007. Neurobehavioral and immunological consequences of prenatal immune activation in rats. Influence of antipsychotics. Neuropsychopharmacology 32:1791–1804. doi: 10.1038/sj.npp.1301292.

23. The PsychENCODE Consortium, Akbarian S, Liu C, Knowles JA, Vaccarino FM, Farnham PJ, Crawford GE, Jaffe AE, Pinto D, Dracheva S, Geschwind DH, Mill J, Nairn AC, Abyzov A, Pochareddy S, Prabhakar S, Weissman S, Sullivan PF, State MW, Weng Z, Peters MA, White KP, Gerstein MB, Senthil G, Lehner T, Sklar P, Sestan N. 2015. The PsychENCODE project. Nature Neuroscience 18:1707–1712. doi: 10.1038/nn.4156.k

24. Zhang Q, Goto H, Akiyoshi-Nishimura S, Prosselkov P, Sano C, Matsukawa H, Yaguchi K, Nakashiba T, Itohara S. 2016. Diversification of behavior and postsynaptic properties by netrin-G presynaptic adhesion family proteins. Molecular Brain 9:e6. doi: 10.1186/s13041-016-0187-5.

25. Zhao M, Kong L, Qu H. 2014. A systems biology approach to identify intelligence quotient score-related genomic regions, and pathways relevant to potential therapeutic treatments. Scientific Reports 4:A4176. doi: 10.1038/srep04176.

26. Johnson MR, Shkura K, Langley SR, Delahaye-Duriez A, Srivastava P, Hill WD, Rackham OJL, Davies G, Harris SE, Moreno-Moral A, Rotival M, Speed D, Petrovski S, Katz A, Hayward C, Porteous DJ, Smith BH, Padmanabhan S, Hocking LJ, Starr JM, Liewald DC, Visconti A, Falchi M, Bottolo L, Rossetti T, Danis B, Mazzuferi M, Foerch P, Grote A, Helmstaedter C, Becker AJ, Kaminski RM, Deary IJ, Petretto E. 2016. Systems genetics identifies a convergent gene network for cognition and neurodevelopmental disease. Nature Neuroscience 19:223–232. doi: 10.1038/nn.4205.

27. Xu K, Schadt EE, Pollard KS, Roussos P, Dudley JT. 2015. Genomic and network patterns of schizophrenia genetic variation in human evolutionary accelerated regions. Molecular Biology and Evolution 32:1148–1160. doi: 10.1093/molbev/msv031.

28. Miller JA, Horvath S, Geschwind DH. 2010. Divergence of human and mouse brain transcriptome highlights Alzheimer disease pathways. Proceedings of the National Academy of Sciences of the United States of America 107:12698–12703. doi: 10.1073/pnas.0914257107.

29. Pollard KS, Salama SR, King B, Kern AD, Dreszer T, Katzman S, Siepel A, Pedersen JS, Bejerano G, Baertsch R, Rosenbloom KR, Kent J, Haussler D. 2006. Forces shaping the fastest evolving regions in the human genome. PLoS Genetics 2:1599–1611. doi: 10.1371/journal.pgen.0020168.

30. Boyd JL, Skove SL, Rouanet JP, Pilaz L-J, Bepler T, Gordân R, Wray GA, Silver DL. 2015. Human-Chimpanzee Differences in a FZD8 Enhancer Alter Cell-Cycle Dynamics in the Developing Neocortex. Current Biology 25:772–779. doi: 10.1016/j.cub.2015.01.041.

31. Zhou H, Hu S, Matveev R, Yu Q, Li J, Khaitovich P, Jin L, Lachmann M, Stoneking M, Fu Q, Tang K. 2015. A Chronological Atlas of Natural Selection in the Human Genome during the Past Half-million Years. BioRxiv. doi: 10.1101/018929.

32. Crabtree GR. 2013. Our fragile intellect. Part I. Trends in Genetics 29:1–3. doi: 10.1016/j.tig.2012.10.002.

33. Hemani G, Shakhbazov K, Westra H-J, Esko T, Henders AK, McRae AF, Yang J, Gibson G, Martin NG, Metspalu A, Franke L, Montgomery GW, Visscher PM, Powell JE. 2014. Detection and replication of epistasis influencing transcription in humans. Nature 508:249–253. doi: 10.1038/nature13005.

34. Jordan DM, Frangakis SG, Golzio C, Cassa CA, Kurtzberg J, Task Force for Neonatal Genomics, Davis EE, Sunyaev SR, Katsanis N. 2015. Identification of cis-suppression of human disease muattions by comparative genomics. Nature 524:225–229. doi: 10.1038/nature14497.

35. Barbosa-Morais NL, Irimia M, Pan Q, Xiong HY, Gueroussov S, Lee LJ, Slobodeniuc V, Kutter C, Watt S, Çolak R, Kim T, Misquitta-Ali CM, Wilson MD, Kim PM, Odom DT, Frey BJ, Blencowe BJ. 2012. The Evolutionary Landscape of Alternative Splicing in Vertebrate Species. Science 338:1587–1593. doi: 10.1126/science.1230612.

36. Lin S, Lin Y, Nery JR, Urich MA, Breschi A, Davis CA, Dobin A, Zaleski C, Beer MA, Chapman WC, Gingeras TR, Ecker JR, Snyder MP. 2014. Comparison of the transcriptional landscapes between human and mouse tissues. Proceedings of the National Academy of Sciences of the United States of America 111:17224–17229. doi: 10.1073/pnas.1413624111.

37. Hyman SE. 2014. How Far Can Mice Carry Autism Research? Cell 158:13–14. doi: 10.1016/j.cell.2014.06.032.

38. Eastwood SL, Harrison PJ. 2008. Decreased mRNA expression of netrin-G1 and netrin-G2 in the temporal lobe in schizophrenia and bipolar disorder. Neuropsychopharmacology 33:933–945. doi: 10.1038/sj.npp.1301457.

39. Aoki-Suzuki M, Yamada K, Meerabux J, Iwayama-Shigeno Y, Ohba H, Iwamoto K, Takao H, Toyota T, Suto Y, Nakatani N, Dean B, Nishimura S, Seki K, Kato T, Itohara S, Nishikawa T, Yoshikawa, T. 2005. A Family-Based Association Study and Gene Expression Analyses of Netrin-G1 and -G2 Genes in Schizophrenia. Biological Psychiatry 57:382–393. doi: 10.1016/j.biopsych.2004.11.022.

40. Nakashiba T, Ikeda T, Nishimura S, Tashiro K, Honjo T, Culotti JG, Itohara, S. 2000. Netrin-G1: a Novel Glycosyl Phosphatidylinositol-Linked Mammalian Netrin That Is Functionally Divergent from Classical Netrins. The Journal of Neuroscience 20:6540–6550.

41. Nakashiba T, Nishimura S, Ikeda T, Itohara S. 2002. Complementary expression and neurite outgrowth activity of netrin-G subfamily members. Mechanisms of Development 111:47–60. doi: 10.1016/S0925-4773(01)00600-1.

42. Song YS, Lee H-J, Prosselkov P, Itohara S, Kim E. 2013. Trans-induced cis interaction in the tripartite NGL-1, netrin-G1 and LAR adhesion complex promotes development of excitatory synapses. Journal of Cell Science 126:4926–4938. doi: 10.1242/jcs.129718.

43. Ule J, Ule A, Spencer J, Williams A, Hu JS, Cline M, Wang H, Clark T, Fraser C, Ruggiu M, Zeeberg BR, Kane D, Weinstein JN, Blume J, Darnell RB. 2005. Nova regulates brain- specific splicing to shape the synapse. Nature Genetics 37:844–852. doi: 10.1038/ng1610.

44. Eastwood SL, Harrison PJ. 2010. Markers of Glutamate Synaptic Transmission and Plasticity Are Increased in the Anterior Cingulate Cortex in Bipolar Disorder. Biological Psychiatry 67:1010–1016. doi: 10.1016/j.biopsych.2009.12.004.

45. Irimia M, Weatheritt RJ, Ellis JD, Parikshak NN, Gonatopoulos-Pournatzis T, Babor M, Quesnel-Vallieres M, Tapial J, Raj B, O'Hanlon D, Barrios-Rodiles M, Sternberg MJE, Cordes SP, Roth FP, Wrana JL, Geschwind DH, Blencowe BJ. 2014. A Highly Conserved Program of Neuronal Microexons Is Misregulated in Autistic Brains. Cell 159:1511–1523. doi: 10.1016/j .cell.2014.11.035.

46. Jung RE and Haier RJ. 2007. The Parieto-Frontal Integration Theory (P-FIT) of intelligence: Converging neuroimaging evidence. Behavioral and Brain Sciences 30:135187. doi: 10.1017/s0140525x07001185.

47. Colom R, Haier RJ, Head K, and Álvarez-Linerac J, Ángeles Quiroga M, Shih PC, Jung RE. 2009. Gray matter correlates of fluid, crystallized, and spatial intelligence: Testing the P-FIT model. Intelligence 37:124–135. doi: 10.1016/j.intell.2008.07.007.

48. Hampshire A, Highfield RR, Parkin BL, Owen AM. 2012. Fractionating Human Intelligence. Neuron 76:1225–1237. doi: 10.1016/j.neuron.2012.06.022.

49. Haier RJ, Karama S, Colom R, Jung R, Johnson W. 2014. A comment on “Fractionating Intelligence” and the peer review process. Intelligence 46:323–332. doi: 10.1016./j.intell.2014.02.007.

50. Song M, Zhou Y, Li J, Liu Y, Tian L, Yu C, Tianzi, J. 2008. Brain spontaneous functional connectivity and intelligence. Neuroimage 41:1168–1176. doi: 10.1016/j.neuroimage.2008.02.036.

51. Glascher J, Tranel D, Paul LK, Rudrauf D, Rorden C, Hornaday A, Grabowski T, Damasio H, Adolphs R. 2009. Lesion Mapping of Cognitive Abilities Linked to Intelligence. Neuron 61:681–691. doi: 10.1016/j.neuron.2009.01.026.

52. Kim J, Ghim J-W, Lee JH, Jung MW. 2013. Neural Correlates of Interval Timing in Rodent Prefrontal Cortex. Journal of Neuroscience 33:13834–13847. doi: 10.1523/jneurosci.1443-13.2013.

53. Markowitz DA, Curtis CE, Pesaran B. 2015. Multiple component networks support working memory in prefrontal cortex. Proceedings of the National Academy of Sciences of the United States of America 112:11084–11089. doi: 10.1073/pnas.1504172112.

54. Spellman T, Rigotti M, Ahmari SE, Fusi S, Gogos JA, Gordon JA. 2015. Hippocampal-prefrontal input supports spatial encoding in working memory. Nature 522:309–314. doi: 10.1038/nature14445.

55. Riga D, Matos MR, Glas A, Smit AB, Spijker S, Van den Oever MC. 2014. Optogenetic dissection of medial prefrontal cortex circuitry. Frontiers in Systems Neuroscience 8:A230. doi: 10.3389/fnsys.2014.00230.

56. Cole MW, Yarkoni T, Repovs G, Anticevic A, Braver TS. 2012. Global Connectivity of Prefrontal Cortex Predicts Cognitive Control and Intelligence. Journal of Neuroscience 32:8988–8999. doi: 10.1523/jneurosci.0536-12.2012.

57. Skeide MA and Friederici AD. 2015. Response to Bornkessel-Schlesewsky *et al*. - towards a nonhuman primate model of language? Trends in Cognitive Sciences. 9:483. doi: 10.1016/j.tics.2015.05.011.

58. Thothathiri M. and Rattinger M. 2015. Ventral and dorsal streams for choosing word order during sentence production. Proceedings of the National Academy of Sciences 112:15456–15461. doi: 10.1073/pnas.1514711112.

59. Gulsuner S and McClellan JM. 2014. *De Novo* Mutations in Schizophrenia Disrupt Genes Co-Expressed in Fetal Prefrontal Cortex. Neuropsychopharmacology 39:238–239. doi: 10.1038/npp.2013.219.

60. Wise SP. 2008. Forward frontal fields: phylogeny and fundamental function. Trends in Neurosciences 31:599–608. doi: 10.1016/j.tins.2008.08.008.

61. Enard W, Przeworski M, Fisher SE, Lai CSL, Wiebe V, Kitano T, Monaco AP, Paabo S. 2002. Molecular evolution of *FOXP2*, a gene involved in speech and language. Nature 418:869–872. doi: 10.1038/nature01025.

62. Krause J, Lalueza-Fox C, Orlando L, Enard W, Green RE, Burbano HA, Hublin J-J, Hänni C, Fortea J, de la Rasilla M, Bertranpetit J, Rosas A, Pääbo S. 2007. The derived *FOXP2* variant of modern humans was shared with neandertals. Current Biology 17:1908–1912. doi: 10.1016/j.cub.2007.10.008.

63. Konopka G, Bomar JM, Winden K, Coppola G, Jonsson ZO, Gao FY, Peng S, Preuss TM, Wohlschlegel JA, Geschwind DH. 2009. Human-specific transcriptional regulation of CNS development genes by FOXP2. Nature 462:213–217. doi: 10.1038/nature08549.

64. Blom N, Gammeltoft S, Brunak S. 1999. Sequence and structure-based prediction of eukaryotic protein phosphorylation sites. Journal of Molecular Biology 294:1351–1362. doi: 10.1006/jmbi.1999.3310.

65. Steentoft C, Vakhrushev SY, Joshi HJ, Kong Y, Vester-Christensen MB, Schjoldager KTBG, Lavrsen K, Dabelsteen S, Pedersen NB, Marcos-Silva L, Gupta R, Bennett EP, Mandel U, Brunak S, Wandall HH, Levery SB, Clausen H. 2013. Precision mapping of the human O-GalNAc glycoproteome through SimpleCell technology. Embo Journal 32:1478–1488. doi: 10.1038/emboj.2013.79.

66. Baenziger JU. 2012. Moving the O-glycoproteome from form to function. Proceedings of the National Academy of Sciences of the United States of America 109:9672–9673. doi: 10.1073/pnas.1206735109.

67. Baenziger JU. 2013. O-mannosylation of cadherins. Proceedings of the National Academy of Sciences of the United States of America 110:20858–20859. doi: 10.1073/pnas.1321827111.

68. Kim S, Burette A. Chung HS, Kwon SK, Woo J, Lee HW, Kim K, Kim H, Weinberg RJ, Kim E. 2006. NGL family PSD-95-interacting adhesion molecules regulate excitatory synapse formation. Nature Neuroscience 9:1294–1301. doi: 10.1038/nn1763.

69. Arbuckle MI, Komiyama NH, Delaney A, Coba M, Garry EM, Rosie R, Allchorne AJ, Forsyth LH, Bence M, Carlisle HJ, O'Dell TJ, Mitchell R, Fleetwood-Walker SM, Grant SGN. 2010. The SH3 domain of postsynaptic density 95 mediates inflammatory pain through phosphatidylinositol-3-kinase recruitment. Embo Reports 11:473–478. doi: 10.1038/embor.2010.63.

70. Soto F, Watkins KL, Johnson RE, Schottler F, Kerschensteiner D. 2013. NGL-2 Regulates Pathway-Specific Neurite Growth and Lamination, Synapse Formation, and Signal Transmission in the Retina. Journal of Neuroscience 33:11949–11959. doi: 10.1523/jneurosci.1521-13.2013.

71. DeNardo LA, de Wit J, Otto-Hitt S, Ghosh A. 2012. NGL-2 Regulates Input-Specific Synapse Development in CA1 Pyramidal Neurons. Neuron 76:762–775. doi: 10.1016/j.neuron.2012.10.013.

72. Abitua PB, Wagner E, Navarrete IA, Levine M. 2012. Identification of a rudimentary neural crest in a non-vertebrate chordate. Nature 492:104–107. doi: 10.1038/nature11589.

73. Green SA, Simoes-Costa M, Bronner ME. 2015. Evolution of vertebrates as viewed from the crest. Nature 520:474–482. doi: 10.1038/nature14436.

74. Stolfi A, Ryan K, Meinertzhagen IA, Christiaen L. 2015. Migratory neuronal progenitors arise from the neural plate borders in tunicates. Nature 527:371–374. doi: 10.1038/nature15758.

75. Abitua PB, Gainous TB, Kaczmarczyk AN, Winchell CJ, Hudson C, Kamata K, Nakagawa M, Tsuda M, Kusakabe, TG, Levine M. 2015. The pre-vertebrate origins of neurogenic placodes. Nature 524:462–465. doi: 10.1038/nature14657.

76. Rada-Iglesias A, Bajpai R, Prescott S, Brugmann SA, Swigut T, Wysocka J. 2012. Epigenomic Annotation of Enhancers Predicts Transcriptional Regulators of Human Neural Crest. Cell Stem Cell 11:1–16. doi: 10.1016/j.stem.2012.07.006.

77. Iskow RC, Gokcumen O, Abyzov A, Malukiewicz J, Zhu Q, Sukumara AT, Pai AA, Mills RE, Habegger L, Cusanovich DA, Rubel MA, Perry GH, Gerstein M, Stone AC, Gilade Y, Lee C. 2012. Regulatory element copy number differences shape primate expression profiles. Proceedings of the National Academy of Sciences of the United States of America. 109:12656–12661. doi: 10.1073/pnas.1205199109.

78. Konopka G, Friedrich T, Davis-Turak J, Winden K, Oldham MC, Gao F, Chen L, Wang G-Z, Luo R, Preuss TM, Geschwind DH. 2012. Human-Specific Transcriptional Networks in the Brain. Neuron 23:601–617. doi: 10.1016/j.neuron.2012.05.034.

79. Liu XL, Somel M, Tang L, Yan Z, Jiang X, Guo S, Yuan Y, He L, Oleksiak A, Zhang Y, Li N, Hu YH, Chen W, Qiu ZL, Pääbo S, Khaitovich P. 2012. Extension of cortical synaptic development distinguishes humans from chimpanzees and macaques. Genome Research 22:611–622. doi: 10.1101/gr.127324.111.

80. McLean CY, Reno PL, Pollen AA, Bassan AI, Capellini TD, Guenther C, Indjeian VB, Lim XH, Menke DB, Schaar BT, Wenger AM, Bejerano G, Kingsley DM. 2011. Human- specific loss of regulatory DNA and the evolution of human-specific traits. Nature 471:216–219. doi: 10.1038/nature09774.

81. Ghalambor CK, Hoke KL, Ruell EW, Fischer EK, Reznick DN, Hughes KA. 2015. Nonadaptive plasticity potentiates rapid adaptive evolution of gene expression in nature. Nature 525:372–375. doi: 10.1038/nature15256.

82. Yaguchi K, Nishimura-Akiyoshi S, Kuroki S, Onodera T, Itohara S. 2014. Identification of transcriptional regulatory elements for *Ntng1* and *Ntng2* genes in mice. Molecular Brain 7:19. doi: 10.1186/1756-6606-7-19.

83. Nishimura-Akiyoshi S, Niimi K, Nakashiba T, and Itohara S. 2007. Axonal netrin-Gs transneuronally determine lamina-specific subdendritic segments. Proceedings of the National Academy of Sciences of the United States of America 104:14801–14806. doi: 10.1073/pnas.0706919104.

84. Matsukawa H, Akiyoshi-Nishimura S, Zhang Q, Lujan R, Yamaguchi K, Goto H, Yaguchi K, Hashikawa T, Sano C, Shigemoto R, Nakashiba T, Itohara S. 2014. NetrinG/NGL Complexes Encode Functional Synaptic Diversification. Journal of Neuroscience 34:15779–15792. doi: 10.1523/jneurosci.1141-14.2014.

85. Zhang F, Shugart YY, Yue W, Cheng Z, Wang G, Zhou Z, Jin C, Yuan J, Liu S, Xu Y. 2015. Increased Variability of Genomic Transcription in Schizophrenia. Scientific Reports 5:e17995. doi: 10.1038/srep17995.

86. Yelin-Bekerman L, Elbaz I, Diber A, Dahary D, Gibbs-Bar L, Alon S, Lerer-Goldshtein T, Appelbaum L. 2015. Hypocretin neuron-specific transcriptome profiling identifies the sleep modulator Kcnh4a. eLife 4:e08638. doi: 10.7554/eLife.08638.

87. McWilliam H, Li W, Uludag M, Squizzato S, Park YM, Buso N, Cowley AP, Lopez R. 2013. Analysis Tool Web Services from the EMBL-EBI. Nucleic Acids Research 41:597–600. doi: 10.1093/nar/gkt376.

88. Voineagu I, Wang X, Johnston P, Lowe JK, Tian Y, Horvath S, Mill J, Cantor RM, Blencowe BJ, Geschwind DH. 2011. Transcriptomic analysis of autistic brain reveals convergent molecular pathology. Nature 474: 380–384. doi: 10.1038/nature10110.

89. O’Roak BJ, Vives L, Girirajan S, Karakoc E, Krumm N, Coe BP, Levy R, Ko A, Lee C, Smith JD, Turner EH, Stanaway IB, Vernot B, Malig M, Baker C, Reilly B, Akey JM, Borenstein E, Rieder MJ, Nickerson DA, Bernier R, Shendure J, Eichler EE. 2012a. Sporadic autism exomes reveal a highly interconnected protein network of *de novo* mutations. Nature 485:246–250. doi: 10.1038/nature10989.

90. O'Roak BJ, Vives L, Fu WQ, Egertson JD, Stanaway IB, Phelps IG, Carvill G, Kumar A, Lee C, Ankenman K, Munson J, Hiatt JB, Turner EH, Levy R, O'Day DR, Krumm N, Coe BP, Martin BK, Borenstein E, Nickerson DA, Mefford HC, Doherty D, Akey JM, Bernier R, Eichler EE, Shendure J. 2012b. Multiplex Targeted Sequencing Identifies Recurrently Mutated Genes in Autism Spectrum Disorders. Science 338:1619–1622. doi: 10.1126/science .1227764.

91. King IF, Yandava CN, Mabb AM, Hsiao JS, Huang HS, Pearson BL, Calabrese JM, Starmer J, Parker JS, Magnuson T, Chamberlain SJ, Philpot BD, Zylka MJ. 2013. Topoisomerases facilitate transcription of long genes linked to autism. Nature 501:58–62. doi: 10.1038/nature12504.

92. Iossifov I, O’Roak BJ, Sanders SJ, Ronemus M, Krumm N, Levy D, Stessman HA, Witherspoon KT, Vives L, Patterson KE, Smith JD, Paeper B, Nickerson DA, Dea J, Dong S, Gonzalez LE, Mandell JD, Mane SM, Murtha MT, Sullivan CA, Walker MF, Waqar Z, Wei L, Willsey AJ, Yamrom B, Lee Y, Grabowska E, Dalkic E, Wang Z, Marks S, Andrews P, Leotta A, Kendall J, Hakker I, Rosenbaum J, Ma B, Rodgers L, Troge J, Narzisi G, Seungtai Y, Schatz MC, Kenny Y, McCombie WR, Shendure J, Eichler EE, State MW, Wigler M. 2014. The contribution of *de novo* coding mutations to autism spectrum disorder. Nature 515:216–221. doi: 10.1038/nature13908.

93. D’Gama AM, Pochareddy S, Li M, Jamuar SS, Reiff RE, Lam A-TN, Sestan N, Walsh CA. 2015. Targeted DNA Sequencing from Autism Spectrum Disorder Brains Implicates Multiple Genetic Mechanisms. Neuron 88:910–917. doi: 10.1016/j.neuron.2015.11.009.

94. Akula N, Barb, J, Jiang, X, Wendland, JR, Choi, KH, Sen, SK, Hou, L, Chen, DTW, Laje, G, Johnson, K, Lipska, BK, Kleinman, JE, Corrada-Bravo, H, Detera-Wadleigh, S, Munson, PJ, McMahon, FJ. 2014. RNA-sequencing of the brain transcriptome implicates dysregulation of neuroplasticity, circadian rhythms and GTPase binding in bipolar disorder. Molecular Psychiatry 19:1179–1185. doi: 10.1038/mp.2013.170.

95. Fukasawa M, Aoki M, Yamada K, Iwayama-Shigeno Y, Takao H, Meerabux J, Toyota T, Nishikawa T, Yoshikawa T. 2004. Case-control association study of human netrin G1 gene in Japanese schizophrenia. Journal of Medical and Dental Sciences 51: 121–128.

96. The Japanese Schizophrenia Sib-Pair Linkage Group (JSSLG), Arinami T, Ohtsuki T, Ishiguro H, Ujike H, Tanaka Y, Morita Y, Mineta M, Takeichi M, Yamada S, Imamura A, Ohara K, Shibuya H, Ohara, K, Suzuki Y, Muratake T, Kaneko N, Someya T, Inada T, Yoshikawa T, Toyota T, Yamada K, Kojima T, Takahashi S, Osamu O, Shinkai T, Nakamura M, Fukuzako H, Hashiguchi T, Niwa S, Ueno T, Tachikawa H, Hori T, Asada T, Nanko S, Kunugi H, Hashimoto R, Ozaki N, Iwata N, Harano M, Arai H, Ohnuma T, Kusumi I, Koyama T, Yoneda H, Fukumaki Y, Shibata H, Kaneko S, Higuchi H, Yasui-Furukori N, Numachi Y, Itokawa M, Okazaki Y. 2005. Genomewide high-density SNP linkage analysis of 236 Japanese families supports the existence of schizophrenia susceptibility loci on chromosomes 1p, 14q, and 20p. American Journal of Human Genetics 77:937–944. doi: 10.1086/498122.

97. Ohtsuki T, Horiuchi Y, Koga M, Ishiguro H, Inada T, Iwata N, Ozaki N, Ujike H, Watanabe Y, Someya T, Arinami T. 2008. Association of polymorphisms in the haplotype block spanning the alternatively spliced exons of the *NTNG1* gene at 1p13.3 with schizophrenia in Japanese populations. Neuroscience Letters 435:194–197. doi: 10.1016/j.neulet.2008.02.053.

98. Zakharyan R, Boyajyan A, Arakelyan A, Gevorgyan A, Mrazek F, Petrek M. 2011. Functional variants of the genes involved in neurodevelopment and susceptibility to schizophrenia in an Armenian population. Human Immunology 72:746–748. doi: 10.1016/j.humimm.2011.05.018.

99. Zhu Y, Yang H, Bi Y, Zhang Y, Zhen C, Xie S, Qin H, He J, Liu L, Liu Y. 2011. Positive association between *NTNG1* and schizophrenia in Chinese Han population. Journal of Genetics 90:499–502.

100. Ayalew M, Le-Niculescu H, Levey DF, Jain N, Changala B, Patel SD, Winiger E, Breier A, Shekhar A, Amdur R, Koller D, Nurnberger JI, Corvin A, Geyer M, Tsuang MT, Salomon D, Schork NJ, Fanous AH, O’Donovan MC, Niculescu AB. 2012. Convergent functional genomics of schizophrenia: from comprehensive understanding to genetic risk prediction. Molecular Psychiatry 17:887–905. doi: 10.1038/mp.2012.37.

101. Wilcox JA and Quadri S. 2014. Replication of NTNG1 association in schizophrenia. Psychiatric Genetics 24:266–268. doi: 10.1097/ypg.0000000000000061.

102. Zhang B, Gaiteri C, Bodea LG, Wang Z, McElwee J, Podtelezhnikov AA, Zhang CS, Xie T, Tran L, Dobrin R, Fluder E, Clurman B, Melquist S, Narayanan M, Suver C, Shah H, Mahajan M, Gillis T, Mysore J, MacDonald ME, Lamb JR, Bennett DA, Molony C, Stone DJ, Gudnason V, Myers AJ, Schadt EE, Neumann H, Zhu J, Emilsson V. 2013. Integrated Systems Approach Identifies Genetic Nodes and Networks in Late-Onset Alzheimer's Disease. Cell 153:707–720. doi: 10.1016/j.cell.2013.03.030.

103. Lagier-Tourenne C, Polymenidou M, Hutt KR, Vu AQ, Baughn M, Huelga SC, Clutario KM, Ling S-C, Liang TY, Mazur C, Wancewicz E, Kim AS, Watt A, Freier S, Hicks GG, Donohue JP, Shiue L, Bennett CF, Ravits J, Cleveland DW, Yeo GW. 2012. Divergent roles of ALS-linked proteins FUS/TLS and TDP-43 intersect in processing long pre- mRNAs. Nature Neuroscience 15:1488–1497. doi: doi:10.1038/nn.3230.

104. Ishigaki S, Masuda A, Fujioka Y, Iguchi Y, Katsuno M, Shibata A, Urano F, Sobue G, Ohno K. 2012. Position-dependent FUS-RNA interactions regulate alternative splicing events and transcriptions. Scientific Reports 2: A529. doi: 10.1038/srep00529.

105. Rogelj B, Easton LE, Bogu GK, Stanton LW, Rot G, Curk T, Zupan B, Sugimoto Y, Modic M, Haberman N, Tollervey J, Fujii R, Takumi T, Shaw CE, Ule J. 2012. Widespread binding of FUS along nascent RNA regulates alternative splicing in the brain. Scientific Reports 2:A603. doi: 10.1038/srep00603.

106. Nakaya T, Alexiou P, Maragkakis M, Chang A, Mourelatos Z. 2013. FUS regulates genes coding for RNA-binding proteins in neurons by binding to their highly conserved introns. RNA 19:498–509. doi: 10.1261/rna.037804.112.

107. Maurano MT, Humbert R, Rynes E, Thurman RE, Haugen E, Wang H, Reynolds AP, Sandstrom R, Qu HZ, Brody J, Shafer A, Neri F, Lee K, Kutyavin T, Stehling-Sun S, Johnson AK, Canfield TK, Giste E, Diegel M, Bates D, Hansen RS, Neph S, Sabo PJ, Heimfeld S, Raubitschek A, Ziegler S, Cotsapas C, Sotoodehnia N, Glass I, Sunyaev SR, Kaul R, Stamatoyannopoulos JA. 2012. Systematic Localization of Common Disease- Associated Variation in Regulatory DNA. Science 337:1190–1195. doi: 10.1126/science .1222794.

108. Wang K, Zhang H, Bloss CS, Duvvuri V, Kaye W, Schork NJ, Berrettini W, Hakonarson H, the Price Foundation Collaborative Group. 2011. A genome-wide association study on common SNPs and rare CNVs in anorexia nervosa. Molecular Psychiatry 16:949–959. doi: 10.1038/mp.2010.107.

109. Boraska V, Franklin CS, Floyd JAB, Thornton LM, Huckins LM, Southam L, Rayner NW, Tachmazidou I, Klump KL, Treasure J, Lewis CM, Schmidt U, Tozzi F, Kiezebrink K, Hebebrand J, Gorwood P, Adan RAH, Kas MJH, Favaro A, Santonastaso P, Fernàndez-Aranda F, Gratacos M, Rybakowsk F, Dmitrzak-Weglarz M, Kaprio J, Keski-Rahkonen A, Raevuori A, Van Furth EF, Slof-Op't Landt MCT, Hudson JI, Reichborn-Kjennerud T, Knudsen GPS, Monteleone P, Kaplan AS, Karwautz A, Hakonarson H, Berrettini WH, Guo Y, Li D, Schork NJ, Komaki G, Ando T, Inoko H, Esko T, Fischer K, Männik K, Metspalu A, Baker JH, Cone RD, Dackor J, DeSocio JE, Hilliard CE, O’Toole JK, Pantel J, Szatkiewicz JP, Taico C, Zerwas S, Trace SE, Davis OSP, Helder S, Bühren K, Burghardt R, de Zwaan M, Egberts K, Ehrlich S, Herpertz-Dahlmann B, Herzog W, Imgart H, Scherag A, Scherag S, Zipfel S, Boni C, Ramoz N, Versini A, Brandys MK, Danner UN, de Kovel C, Hendriks J, Koeleman BPC, Ophoff RA, Strengman E, van Elburg AA, Bruson A, Clementi M, Degortes D, Forzan M, Tenconi E, Docampo E, Escaramis G, Jimenez-Murcia S, Lissowska J, Rajewski A, Szeszenia-Dabrowska N, Slopien A, Hauser J, Karhunen L, Meulenbelt I, Slagboom PE, Tortorella A, Maj M, Dedoussis G, Dikeos D, Gonidakis F, Tziouvas K, Tsitsika A, Papezova H, Slachtova L, Martaskova D, Kennedy JL, Levitan RD, Yilmaz Z, Huemer J, Koubek D, Merl E, Wagner G, Lichtenstein P, Breen G, Cohen-Woods S, Farmer A, McGuffin P, Cichon S, Giegling I, Herms S, Rujescu D, Schreiber S, Wichmann H-E, Dina C, Sladek R, Gambaro G, Soranzo N, Julia A, Marsal S, Rabionet R, Gaborieau V, Dick DM, Palotie A, Ripatti S, Widen E, Andreassen OA, Espeseth T, Lundervold A, Reinvang I, Steen VM, Le Hellard S, Mattingsdal M, Ntalla I, Bencko V, Foretova L, Janout V, Navratilova M, Gallinger S, Pinto D, Scherer SW, Aschauer H, Carlberg L, Schosser A, Alfredsson L, Ding B, Klareskog L, Padyukov L, Courtet P, Guillaume S, Jaussent I, Finan C, Kalsi G, Roberts M, Logan DW, Peltonen L, Ritchie GRS, Barrett JC, The Wellcome Trust Case Control Consortium 3, Estivill X, Hinney A, Sullivan PF, Collier DA, Zeggini E, Bulik CM. 2014. A genome-wide association study of anorexia nervosa. Molecular Psychiatry 19:1084–1094. doi: 1085-1094.10.1038/mp.2013.187.

110. van Kuilenburg ABP, Meijer J, Mul ANPM, Hennekam RCM, Hoovers JMN, de Die-Smulders CEM, Weber P, Mori AC, Bierau J., Fowler B, Macke K, Sass JO, Meinsma R, Hennermann JB, Miny P, Zoetekouw L, Vijzelaar R, Nicolai J, Ylstra, B, Rubio-Gozalbo ME. 2009. Analysis of severely affected patients with dihydropyrimidine dehydrogenase deficiency reveals large intragenic rearrangements of *DPYD* and a de novo interstitial deletion del(1)(p13.3p21.3). Human Genetics 125:581–590. doi: 10.1007/s00439-009-0653-6

111. Gilissen C, Hehir-Kwa JY, Thung DT, van de Vorst M, van Bon BWM, Willemsen MH, Kwint M, Janssen IM, Hoischen A, Schenck A, Leach R, Klein R, Tearle R, Bo T, Pfundt R, Yntema HG, de Vries BBA, Kleefstra T, Brunner HG, Vissers LELM, Veltman JA. 2014. Genome sequencing identifies major causes of severe intellectual disability. Nature 511:344–347. doi: 10.1038/nature13394.

112. Stepanyan A, Zakharyan R, Boyajyan A. 2013. The netrin G1 gene rs628117 polymorphism is associated with ischemic stroke. Neuroscience Letters 549:74–77. doi: 10.1016/j.neulet.2013.05.066.

113. Bisgaard A-M, Rasmussen LN, Moller HU, Kirchhoff M, Bryndorf T. 2007. Interstitial deletion of the short arm of chromosome 1 (1p13.1p21.1) in a girl with mental retardation, short stature and colobomata. Clinical Dysmorphology 16:109–112. doi: 10.1097/01.mcd.0000228425.89660.bf.

114. Stewart SE, Yu D, Scharf JM, Neale BM, Fagerness JA, Mathews CA, Arnold PD, Evans PD, Gamazon ER, Osiecki L, McGrath L, Haddad S, Crane J, Hezel D, Illman C, Mayerfeld C, Konkashbaev A, Liu C, Pluzhnikov A, Tikhomirov A, Edlund CK, Rauch SL, Moessner R, Falkai P, Maier W, Ruhrmann S, Grabe HJ, Lennertz L, Wagner M, Bellodi L, Cavallini MC, Richter MA, Cook EH, Kennedy JL, Rosenberg D, Stein DJ, Hemmings SMJ, Lochner C, Azzam A, Chavira DA, Fournier E, Garrido H, Sheppard B, Umana P, Murphy DL, Wendland JR, Veenstra-VanderWeele J, Denys D, Blom R, Deforce D, Van Nieuwerburgh F, Westenberg HGM, Walitza S, Egberts K, Renner T, Miguel EC, Cappi C, Hounie AG, do Rosario MC, Sampaio AS, Vallada H, Nicolini H, Lanzagorta N, Camarena B, Delorme R, Leboyer M, Pato CN, Pato MT, Voyiaziakis E, Heutink P, Cath DC, Posthuma D, Smit JH, Samuels J, Bienvenu OJ, Cullen B, Fyer AJ, Grados MA, Greenberg BD, McCracken JT, Riddle MA, Wang Y, Coric V, Leckman JF, Bloch M, Pittenger C, Eapen V, Black DW, Ophoff RA, Strengman E, Cusi D, Turiel M, Frau F, Macciardi F, Gibbs JR, Cookson MR, Singleton A for the North American Brain Expression Consorium, Hardy J for the UK Human Brain Expression Database, Crenshaw AT, Parkin MA, Mirel DB, Conti DV, Purcell S, Nestadt G, Hanna GL, Jenike MA, Knowles JA, Cox N, Pauls DL. 2013. Genome-wide association study of obsessive- compulsive disorder. Molecular Psychiatry 18:788–798. doi: 10.1038/mp.2012.85.

115. Lesnick TG, Papapetropoulos S, Mash DC, Ffrench-Mullen J, Shehadeh L, de Andrade M, Henley JR, Rocca WA, Ahlskog JE, Maraganore DM. 2007. A genomic pathway approach to a complex disease: Axon guidance and Parkinson disease. PLoS Genetics 3:984–995. doi: 10.1371/journal.pgen.0030098.

116. Borg I, Freude K, Kubart SK, Hoffmann K, Menzel C, Laccone F, Firth H, FergusonSmith MA, Tommerup N, Ropers HH, Sargan D, Kalscheuer VM. 2005. Disruption of Netrin G1 by a balanced chromosome translocation in a girl with Rett syndrome. European Journal of Human Genetics 13:921–927. doi: 10.1038/sj.ejhg.5201429.

117. Archer HL, Evans JC, Millar DS, Thompson PW, Kerr AM, Leonard H, Christodoulou J, Ravine D, Lazarou L, Grove L, Verity C, Whatley SD, Pitz DT, Sampson JR, Clarke AJ. 2006. *NTNG1* mutations are a rare cause of Rett syndrome. American Journal of Medical Genetics Part A 140A:691–694. doi: 10.1002/ajmg.a.31133.

118. Nectoux J, Girard B, Bahi-Buisson N, Prieur F, Afenjar A, Rosas-Vargas H, Chelly J, Bienvenu T. 2007. *Netrin G1* mutations are an uncommon cause of atypical Rett syndrome with or without epilepsy. Pediatric Neurology 37:270–274. doi: 10.1016/j.pediatrneurol.2007.06.002.

119. Pan Y, Fang M, Shen L, Wang L, Lv Y, Xi Z, Chen D, Li C, Wang X. 2010. Abnormal expression of netrin-G2 in temporal lobe epilepsy neurons in humans and a rat model. Experimental Neurology 224:340–346. doi: 10.1016/j.expneurol.2010.04.001.

120. Scharf JM, Yu D, Mathews CA, Neale BM, Stewart SE, Fagerness JA, Evans P, Gamazon E, Edlund CK, Service SK, Tikhomirov A, Osiecki L, Illmann C, Pluzhnikov A, Konkashbaev A, Davis LK, Han B, Crane J, Moorjani P, Crenshaw AT, Parkin MA, Reus VI, Lowe TL, Rangel-Lugo M, Chouinard S, Dion Y, Girard S, Cath DC, Smit JH, King RA, Fernandez TV, Leckman JF, Kidd KK, Kidd JR, Pakstis AJ, State MW, Herrera LD, Romero R, Fournier E, Sandor P, Barr CL, Phan N, Gross-Tsur V, Benarroch F, Pollak Y, Budman CL, Bruun RD, Erenberg G, Naarden AL, Lee PC, Weiss N, Kremeyer B, Berrio GB, Campbell DD, Silgado JCC, Ochoa WC, Restrepo SCM, Muller H, Duarte AVV, Lyon GJ, Leppert M, Morgan J, Weiss R, Grados MA, Anderson K, Davarya S, Singer H, Walkup J, Jankovic J, Tischfield JA, Heiman GA, Gilbert DL, Hoekstra PJ, Robertson MM, Kurlan R, Liu C, Gibbs JR, Singleton A for the North American Brain Expression Consorium, Hardy J for the UK Human Brain Expression Database, Strengman E, Ophoff RA, Wagner M, Moessner R, Mirel DB, Posthuma D, Sabatti C, Eskin E, Conti DV, Knowles JA, Ruiz-Linares A, Rouleau GA, Purcell S, Heutink P, Oostra BA, McMahon WM, Freimer NB, Cox NJ, Pauls DL. 2013. Genome-wide association study of Tourette's syndrome. Molecular Psychiatry 18:721–728. doi: 10.1038/mp.2012.69.

121. del Rosario RCH, Rayan NA, Prabhakar S. 2014. Noncoding Origins of Anthropoid Traits and a New Null Model of Transposon Functionalization. Genome Research 24:1469–1484. doi: 10.1101/gr.168963.113.

122. Petersen TN, Brunak S, von Heijne G, Nielsen H. 2011. SignalP 4.0: discriminating signal peptides from transmembrane regions. Nature Methods 8:785–786. doi: 10.1038/nmeth.1701.

123. Eisenhaber B, Bork P, Yuan YP, Loffler G, Eisenhaber F. 2000. Automated annotation of GPI anchor sites: case study *C. elegans*. Trends in Biochemical Sciences 25:340–341. doi: 10.1016/s0968-0004(00)01601-7.

124. Prabhakar S, Noonan JP, Pääbo S, Rubin EM. 2006. Accelerated evolution of conserved noncoding sequences in humans. Science 314:786–786. doi: 10.1126/science.1130738.

125. Meerabux JMA, Ohba H, Fukasawa M, Suto Y, Aoki-Suzuki M, Nakashiba T, Nishimura S, Itohara S, Yoshikawa T. 2005. Human netrin-G1 isoforms show evidence of differential expression. Genomics 86:112–116. doi: 10.1016/j.ygeno.2005.04.004.

